# DNA writing at a single genomic site enables lineage tracing and analog recording in mammalian cells

**DOI:** 10.1101/639120

**Authors:** Theresa B. Loveless, Joseph H. Grotts, Mason W. Schechter, Elmira Forouzmand, Courtney K. Carlson, Bijan S. Agahi, Guohao Liang, Michelle Ficht, Beide Liu, Xiaohui Xie, Chang C. Liu

## Abstract

The study of intricate cellular and developmental processes in the context of complex multicellular organisms is difficult because it can require the non-destructive observation of thousands, millions, or even billions of cells deep within an animal. To address this difficulty, several groups have recently reported CRISPR-based DNA recorders that convert transient cellular experiences and processes into changes in the genome, which can then be read by sequencing in high-throughput. However, existing DNA recorders act primarily by *erasing* DNA: they use the random accumulation of CRISPR-induced deletions to record information. This is problematic because in the limit of progressive deletion, no record remains. Here, we present a new type of DNA recorder that acts primarily by *writing* new DNA. Our system, called CHYRON (Cell HistorY Recording by Ordered iNsertion), inserts random nucleotides at a single locus in temporal order *in vivo* and can be applied as an evolving lineage tracer as well as a recorder of user-selected cellular stimuli. As a lineage tracer, CHYRON allowed us to perfectly reconstruct the population lineage relationships among 16 groups of human cells descended from four starting groups that were subject to a series of splitting steps. In this experiment, CHYRON progressively wrote and retained base insertions in 20% percent of cells where the average amount written was 8.4 bp (~14.5 bits), reflecting high information content and density. As a stimulus recorder, we showed that when the CHYRON machinery was placed under the control of a stress-responsive promoter, the frequency and length of writing reflected the dose and duration of the stress. We believe CHYRON represents a conceptual advance in DNA recording technologies where writing rather than erasing becomes the primary mode of information accumulation. With further engineering of CHYRON’s components to increase writing efficiency, CHYRON should lead to single-cell-resolution recording of lineage and other information through long periods of time in complex animals or tumors, advancing the pursuit of a full picture of mammalian development.

## Introduction

Non-destructive observation of living organisms is a cornerstone of biology. Over time, our ability to observe ever-smaller organisms, individual cells within multicellular organisms, and molecules within cells has improved progressively with advances in microscopy and the continuing development of genetically-encoded labels that can be imaged non-destructively *(e.g.,* GFP). However, live imaging of single cells in intact organisms is still severely constrained by context and scale. Animals, for example, tend to be opaque and even when developmental processes are accessible to microscopy^1^, cell tracking poses significant computational and data management challenges when the number of cells reaches only tens of thousands^1^. An alternative paradigm to non-destructive direct observation is DNA recording. In DNA recording, transient cellular events are engineered to trigger permanent mutations in a cell’s own genome (**Figure 1A-B**). Since DNA is both durable and propagating, and since the throughput of DNA sequencing is in the hundreds of millions of unique DNA molecules, the long-term behavior of cells can be stored as mutations in DNA and read out later at unprecedented depth. Although the reading step is destructive, recording is not, creating an effective alternative to direct observation that can scale to millions of cells in opaque model animals such as mice.

**Figure 1.**
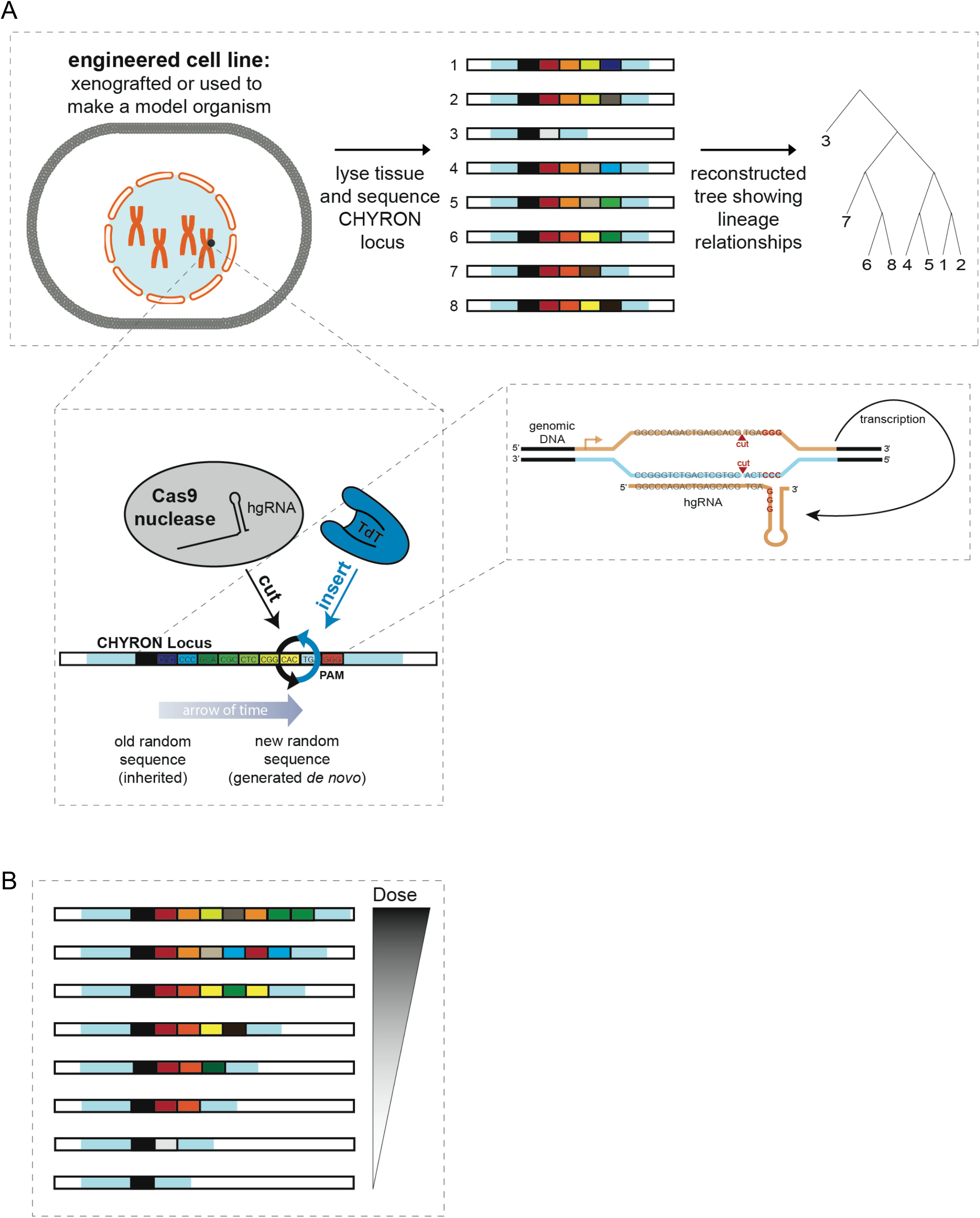
Design of the CHYRON locus. **(A)** The constitutive expression of Cas9 and TdT mediates the ordered acquisition of random insertion mutations, represented as differently-colored boxes, written by TdT. Each cell is represented by one unique sequence, which can be compared to other sequences to reconstruct the lineage of the cells. The CHYRON locus encodes a homing or self-targeting guide RNA, which directs Cas9 to the DNA that encodes it^3–4^. This architecture allows multiple rounds of cutting by Cas9 and writing by TdT. **(B)** If Cas9 and TdT expression is tied to a stimulus, the lengths of the TdT-mediated insertions can be interpreted to determine each cell’s relative dose of the stimulus.

This paradigm of DNA recording has recently seen a transformation with the development of genetically-encoded CRISPR-based systems that drive rapid mutational accumulation at neutral loci in a cell’s genome^2–16^. When the activity of such systems is linked to the presence of an arbitrary biological stimulus, accumulated mutations become a record of the strength and duration of exposure to the stimulus^3,5,7^; and when activity is constitutive, accumulated mutations capture lineage relationships among individual cells^2,4–6,11–14^. Two recent architectures for CRISPR-based recording systems are particularly amenable to recording in extremely large numbers of mammalian cells. The first architecture relies on arrays of *Streptococcus pyogenes* Cas9-target sites^2,12,14,17^. Here, Cas9 targets random elements of the array to generate insertions or deletions (indels) at array elements. The progressive accumulation of indels across the array yields a mutant array in each cell that can be used to infer lineage relationships or a cell’s history of exposure to a stimulus. The second architecture relies on a self-targeting^3^ or homing guide RNA^4^ (hgRNA) that directs Cas9 to the very locus from which the hgRNA is expressed (**Figure 1C**). Here, the hgRNA locus changes over time to generate a pattern of indels at the locus that reflects lineage information or exposure to stimuli. These two types of systems have been used to identify the early and late embryonic origin of thousands of cell lineages in adult zebrafish^2,6,13^, to study embryogenesis in mice^12^, and to record inflammation exposure in 293T cells implanted into mice treated with lipopolysaccharide^3^. However, these systems suffer from a common and critical problem: the predominant repair outcomes of CRISPR nuclease-induced double-strand breaks (DSBs) in mammalian cells are deletions^18^. Since the progressive accumulation of deletions at a single site will quickly corrupt or remove previous deletions^3,4^, and continuous editing of an array of sites at a single locus can lead to multiple simultaneous cuts and the loss of intervening information^2^, these DNA recording systems are limited in their information capacity and durability. In other words, existing DNA recording systems *erase* DNA as their primary mode of recording information. Although patterns of erasures contain new information, the inherent contradiction in removing DNA to add information presents fundamental challenges in the continued development of existing DNA recorder designs.

An ideal DNA recorder should instead be able to *write* DNA. We present such a recorder by constructing a mutating DNA barcode called CHYRON (Cell HistorY Recording by Ordered iNsertion) (**Figure 1A-B**). CHYRON combines a Cas9 nuclease with an hgRNA and a DNA polymerase, terminal deoxynucleotidyl transferase (TdT)^19,20^, capable of efficiently writing random nucleotides (nts) at Cas9-induced DSBs. These newly-written nts are then incorporated into the repaired DSB to produce a durable insertion mutation consisting of random base pairs (bps). Since the hgRNA repeatedly directs Cas9 to cut its own locus at a defined location relative to the PAM^3,4^, cycles of cutting, writing by TdT, and repair cause continuous and ordered insertional mutagenesis (**Figure 1A**). We describe the successful implementation of CHYRON and apply it to lineage reconstruction and recording of hypoxia. We find that the information generated at a single <100 bp CHYRON locus is sufficient to reconstruct the relatedness of populations containing thousands of lineages as well as to report on the duration and dose of exposure to a hypoxia mimic. This opens up the possibility of following mammalian development with unprecedented ease and depth, with the ability to distinguish the hundreds of millions of cells produced during mammalian development, or profiling the heterogeneous responses of each cell in a population to unevenly-distributed or dynamic stresses.

## Results and Discussion

### TdT mediates random insertional mutagenesis at Cas9 cut sites

The core functionality of CHYRON relies on insertional mutagenesis by TdT. Therefore, we first tested whether a single round of Cas9 cutting and DSB repair could be intercepted by TdT to generate insertions. In ~10^5^ HEK293T cells, we targeted Cas9 to a genomic locus with or without the ectopic expression of TdT. We then analyzed repair outcomes by PCR amplification of the target locus followed by next-generation sequencing (NGS), taking care to capture substitution, deletion, and insertion mutations equally (see **Methods**). We found that without TdT, the dominant mutation present at the Cas9 target site was a deletion (84%) or 1 bp insertion (13%) (**Figure 2A**), consistent with previous literature^18,21^. However, with TdT, the dominant mutations were insertions (74%) (**Figure 2A**), with an average length of 2.8 bp (**Figure 2B**). This dramatic shift in repair outcomes in a single round of editing suggested to us that once combined with an hgRNA, CHYRON should be able to write new DNA over multiple rounds with only moderate sequence corruption or loss through deletions.

**Figure 2.**
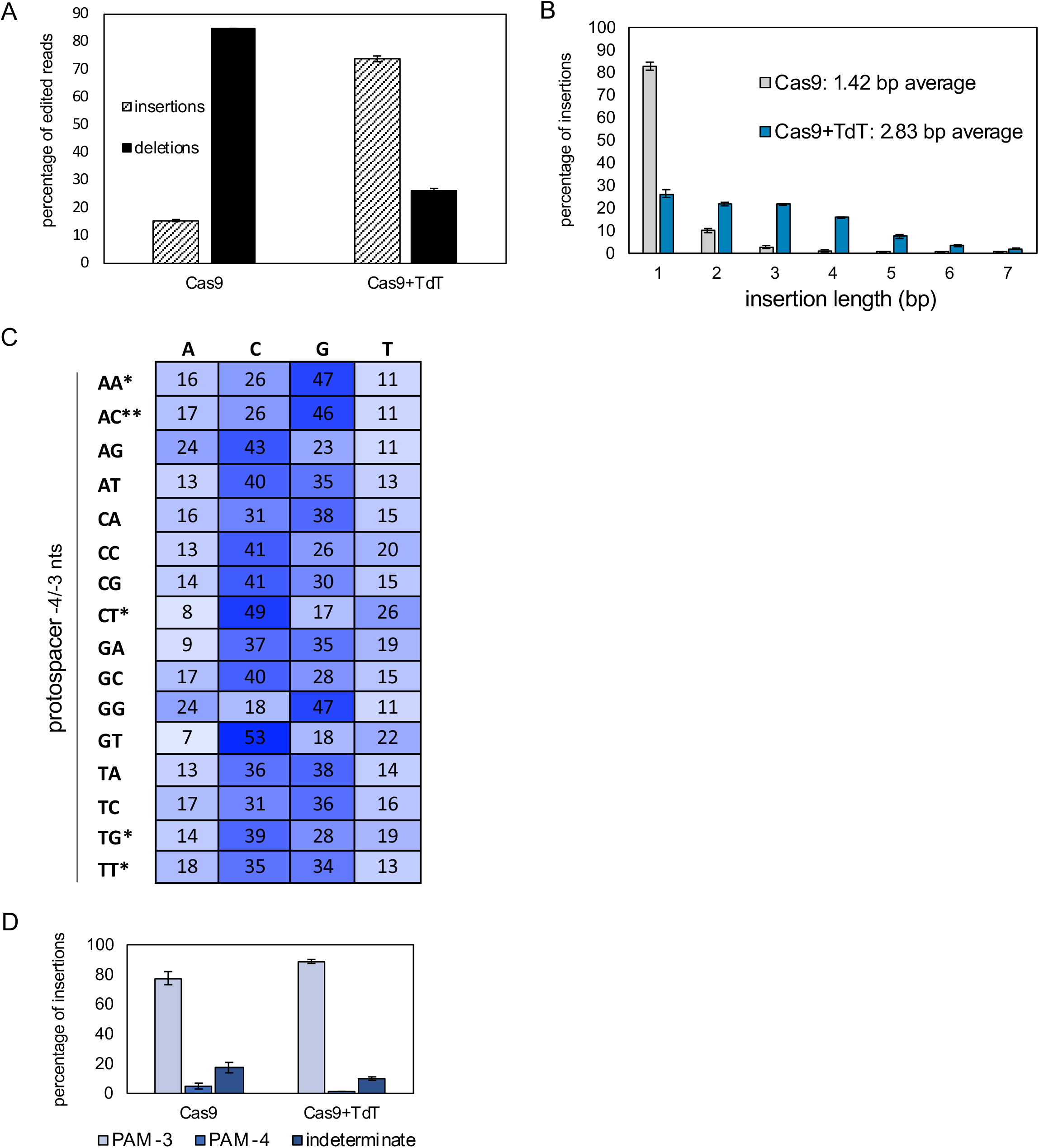
TdT writes DNA at a Cas9-induced DSB. **(A)** Expression of TdT promoted insertion mutations. 293T cells were transfected with plasmids expressing Cas9 and TdT, or Cas9 alone, and an sgRNA against a genomic site (HEK293site3^37^). Three days later, cells were collected, DNA was extracted, and the targeted genomic site was amplified by PCR and sequenced by NGS. Bars represent the mean of six replicates (two technical replicates each of three biological replicates). Error bars = ± stdev. Sequences were placed in one of three categories: unchanged, pure insertions (insertions), or any sequence that leads to a loss of information (deletions). “Deletions” included pure deletions; mixtures of insertion, deletion, and substitution mutations; and pure substitutions (2% of all edited sequences). **(B)** Expression of TdT resulted in longer insertion mutations than those minority insertions created in the presence of Cas9 alone, suggesting that TdT acts as a DNA writer. Of the pool of pure insertions, the percentage of each length was calculated and plotted. Bars represent the mean of the six replicates. Error bars = ±stdev. **(C)** Insertion sequences generated by TdT were random, but had a bias toward G and C nucleotides. 293T cells were transfected with plasmids expressing Cas9 and TdT, and one of 16 sgRNAs against different genomic sites. The target protospacers were chosen to have all possible combinations of nucleotides at the sites 4 and 3 nt upstream of the PAM sequence on the top (nontarget) strand. The proportions of each nucleotide (on the top strand) found in all pure insertion sequences 4 bp in length were calculated for each protospacer. Data shown are the average of four replicates (two technical replicates each of two biological replicates), except those marked with *, which are the average of two technical replicates of a single biological replicate, and the row marked with **, which are the average of two biological replicates. **(D)** TdT-mediated insertions were added 3 bp upstream of the PAM. For the pure insertions shown in (A), the position of the insertion was determined if the insertion sequence made this determination possible. Insertions were annotated as having an “indeterminate” position, for example, if the 3’ nt of the insertion was identical to the protospacer nt 5’ of where the insertion was placed.

TdT-mediated insertions must have a wide diversity of possible sequences for CHYRON to be an effective DNA recorder. By analyzing 16 different target sites, we found that TdT-mediated insertional mutagenesis generated an average insertion length of 2.9 bp (5.2 bits) per round (**Table S1**). To calculate these values, we characterized the single-round length of insertions at Cas9 targets in the presence of TdT. As shown in **Figure 2B**, insertions were commonly 1-4 bp, with some longer insertions that raised the mean length to 2.9 bp. To measure biases in the inserted bps in order to determine the average number of bits of information in each bp written, we designed single guide RNAs (sgRNAs) to recruit Cas9 to a panel of genomic sites. Our choice of target sites was made to explore all 16 possible pairs of the −4 and −3 nts relative to the PAM, since Cas9 canonically creates a blunt cut between these −4 and – 3 positions and there is a known influence of these nts on editing outcomes^22–24^. We found that single-bp insertion outcomes were likeliest to have the identity of the −4 nt, a bias that was independent of TdT (**Table S1**) and that could be explained by staggered Cas9 cuts (−4 relative to the PAM in the nontemplate strand and +3 relative to PAM in the template strand)^22–24^ or by the requirement for cohesive ends in order to promote re-ligation of a DSB. However, in longer insertions, we found that the identity bias of inserted bps was driven by TdT, which is known to prefer the insertion of Gs *in vitro* and during VDJ recombination^25^. This resulted in our overall observation that Gs and Cs comprised ~70% of nts inserted (**Figure 2C** and **Table S1**). Our results also indicated that TdT added to both 3’ ends of the DSB, since we saw a similar preference for both Gs and Cs in longer insertions. However, some target sites resulted in insertions with a strong bias for G over C or vice versa, suggesting that TdT preferred to add nts to one DNA terminus at a DSB over the other in a sequence-dependent manner. To determine the information content of TdT-mediated insertions, we calculated the entropy per bp for each insertion length at all 16 genomic target sites we tested (**Table S1**). (This calculation implies that the biases of bps inserted were independent of each other, which is a reasonable approximation of what we observe – see Table S6.) From these entropies per bp, we calculated the expected value for entropy per round of editing at each site. Averaging these expected values over all 16 sites gives 5.2 bits from a mean insertion length of 2.9 bp. Therefore, TdT-mediated insertions encode an average of 1.78 bits of information per bp (**Table S1**), compared to 2 bits if all 4 bases were added randomly.

### CHYRON_20_ accumulates ordered insertion mutations in multiple rounds

A recording locus should autonomously accumulate mutations over multiple rounds of activity so that cellular and developmental processes occurring over time can be captured. To achieve multiple rounds of DNA writing, we combined TdT with an hgRNA locus^3,4^ to establish CHYRON. Because Cas9-induced DSBs are consistently generated between the −4 and −3 nts relative to the PAM of the hgRNA locus (**Figure 2D**), rounds of TdT-mediated insertion mutations should follow in order when repeated. This makes CHYRON an ideal recording locus because (1) new insertions will neither remove nor corrupt previous insertions and (2) insertions are directionally arranged in the exact order in which they are added, simplifying inference of lineage from the mutational information recorded.

To demonstrate repeated and ordered insertional mutagenesis, we integrated an hgRNA locus, including a 20-nt spacer, at a single site in 293T cells (**Figure 3A**). We call this locus CHYRON_20_, using the subscript to distinguish this specific instantiation of CHYRON with a 20-nt hgRNA spacer from others discussed later. When the cell line containing CHYRON_20_ was transfected with a plasmid expressing Cas9 and TdT for three days, insertions accumulated at the locus as expected (**Figure 3B-C**). (For simplicity, we will refer to hgRNA loci as CHYRON loci when in the presence of TdT, and to the cell line bearing the integrated CHYRON_20_ locus as 293T-CHYRON_20_.) As a comparison, we carried out a similar experiment where a genomic locus with the same spacer sequence as CHYRON_20_ was targeted by an sgRNA, thereby allowing for only a single round of editing (**Figure 3D**). We found that 1-2 bp insertions were less abundant and longer insertions were more abundant at the CHYRON_20_ locus compared to the genomic locus targeted by an sgRNA. This difference strongly suggested that the CHYRON_20_ locus was edited in multiple successive rounds. In order to show multi-round editing conclusively, we isolated 293T-CHYRON_20_ cells that had acquired an insertion at the CHYRON_20_ locus to near-clonality, then transfected them with Cas9 and TdT again. Although further editing was inefficient, new insertions were abundantly observed and all new insertions were found precisely downstream of the original insertion (**Figure 3E-F** and **Figure S3B-C**).

**Figure 3.**
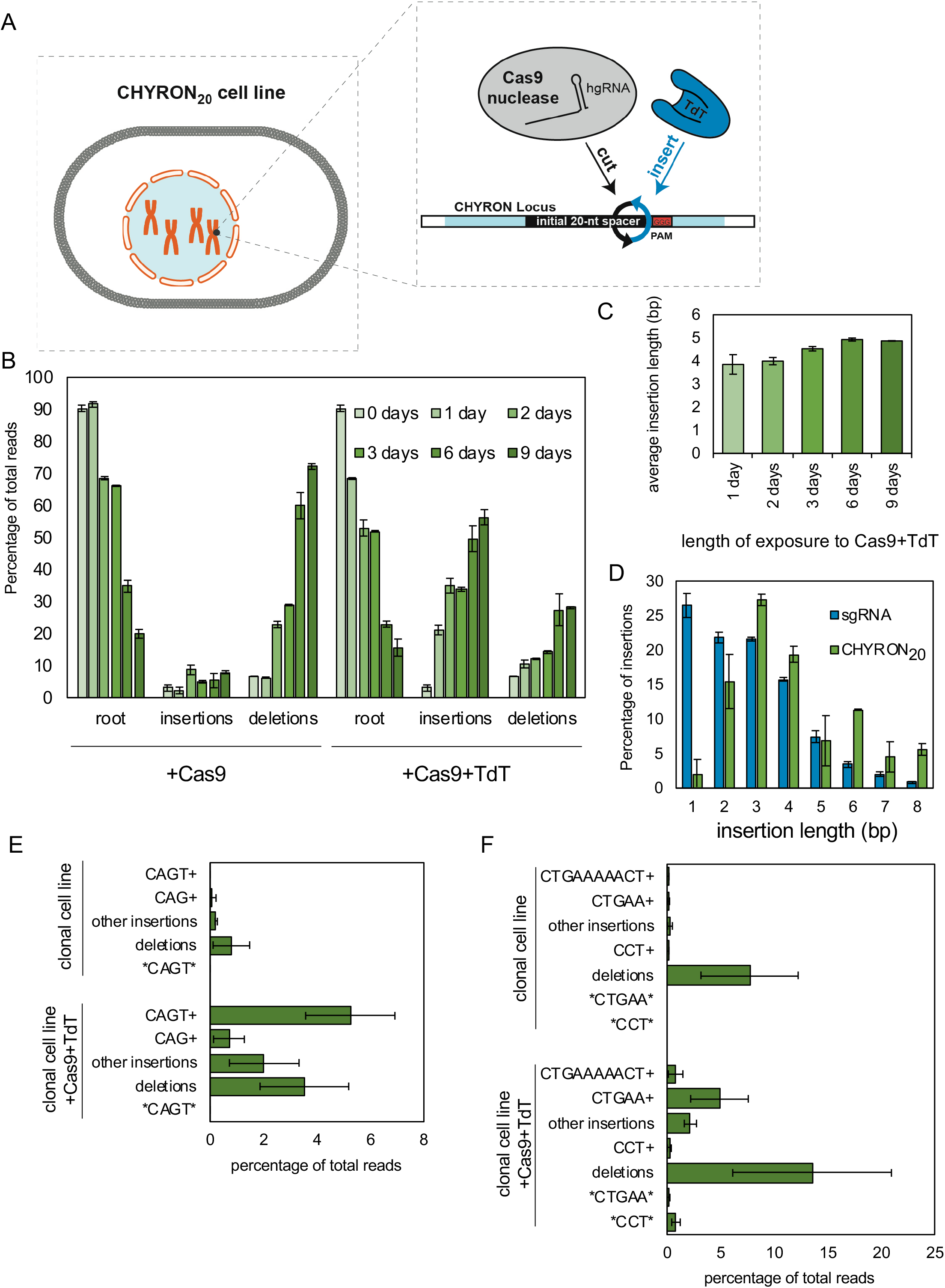
An integrated hgRNA accumulates insertions in multiple rounds. **(A)** A clonal 293T cell line bearing an integrated hgRNA (hereafter 293T-CHYRON_20_) was created so that expression of Cas9 and TdT resulted in multiple rounds of insertion of random nucleotides in an ordered fashion. **(B)** 293T-CHYRON_20_ cells accumulated insertions and deletions over time when exposed to Cas9 and TdT. 293T-CHYRON_20_ cells were transfected with a plasmid expressing Cas9 and, optionally, TdT for the indicated time before collection. Cells were re-transfected every three days. The hgRNA locus was sequenced and each sequence was annotated as unchanged (root), pure insertion (insertions), or any sequence that involves a loss of information (deletions). Bars represent the average of two technical replicates. Error bars= ± stdev. **(C)** Insertions grew longer, on average, over time, until the six day timepoint, then stopped growing. The average length of pure insertions at each timepoint was calculated. **(D)** Longer insertions were more abundant for an hgRNA than for a protospacer targeted in a single round. Insertion lengths at the CHYRON_20_ locus after 3 days of Cas9 and TdT expression were compared to insertions at a genomic site with the same spacer sequence targeted with an sgRNA (data from Figure 2B). **(E)** Cas9 and TdT mediated multiple rounds of editing on an integrated hgRNA. The 293T-CHYRON_20_ cell line was transfected with Cas9 and TdT to induce insertions, then purified to near-clonality. This near-clonal cell line bearing an insertion with the sequence CAGT was then transfected again with a plasmid expressing Cas9 and TdT. These cells, and an untransfected control, were grown for 6-15 days, then collected. The CHYRON locus was sequenced and editing outcomes were determined to be the root CHYRON_20_ sequence (root), deletions, a CAGT insertion (CAGT), an insertion containing the prefix CAGT or CAG (CAGT+ or CAG+, respectively), an insertion containing the sequence CAGT other than as a prefix (*CAGT*), or other insertions. Error bars=±stdev of two technical replicates each of the 6 and 15 day timepoints. A plot showing the most-abundant root and CAGT sequences can be found in Figure S3B. **(F)** Another near-clonal cell line was isolated, this one bearing the insertions CTGAAAAACT and CCT. This cell line was treated and sequenced as in (D) and editing outcomes were determined to be the root CHYRON_20_ sequence (root), deletions, both isolated insertions (CTGAAAAACT and CCT), an insertion containing these insertions or a shortened version as a prefix (CTGAAAAACT+, CCT+, CTGAA+), an insertion containing the sequences CTGAA or CCT other than as a prefix (*CTGAA* and *CCT*), or other insertions. Error bars=±stdev of two technical replicates each of the 6 and 15 day timepoints. A plot showing the most-abundant root, CTGAAAAACT, and CCT sequences can be found in Figure S3C.

CHYRON_20_ gave our basic desired behavior, progressively generating 8.1 bits of information on average via ordered insertions of short random bp stretches, but it was clear CHYRON_20_ could be improved. For example, we deduced that CHYRON_20_ only underwent approximately two rounds of editing, because the proportion of deletion-containing sequences was ~35% of the converted CHYRON_20_ loci and we knew that a single round of editing generates ~26% deletions (**Figure 2A**), requiring two rounds to give ~35%. We also found that when 293T-CHYRON_20_ cells were transfected with Cas9 and TdT over 9 days, the average insertion length plateaued at 4.9 bp, which was already reached after 6 days of Cas9/TdT expression (**Figure 3C**). This was shorter than the average 5.8 bp length that we expected for two rounds, with the discrepancy suggesting that shorter initial insertions were disproportionately likely to continue to edit. Therefore, we sought to improve CHYRON_20_ to be capable of more rounds of activity, which should result in a greater potential diversity of insertion sequences.

### CHYRON_16i_ accumulates an average of 8.4 inserted bps over an average of three rounds

The failure of 293T-CHYRON_20_ cells to write more than an average of 4.9 bp has two likely explanations: silencing of the CHYRON locus or reduced efficiency of the hgRNA through increased length and/or secondary structure associated with continued rounds of TdT-mediated insertions. To address these potential problems, we created two new cell lines (**Figure 4A**), 293T-CHYRON_20i_ and 293T-CHYRON_16i_, both of which have the CHYRON locus flanked by chromatin insulator sequences^26^ and integrated at the *AAVS1* safe harbor locus in 293T cells. CHYRON_20i_ starts with a 20 nt spacer, which operates at peak activity, while CHYRON_16i_ starts with a 16 nt spacer, which is predicted to have very low activity initially^27^, but has more room to accumulate insertions before it reaches lengths that prohibitively reduce editing efficiency^3,4^. These new CHYRONs significantly outperformed the original CHYRON_20_. Specifically, 293T-CHYRON_20i_ cells, in contrast to 293T-CHYRON_20_ cells, continued to accumulate longer insertions throughout the entire 9-day time course and resulted in CHYRON_20i_ loci that reached a final length of 5.7 generated bps on average (**Figure 4B-C**); and 293T-CHYRON_16i_ cells, even though they had a lower overall editing efficiency due to the low starting activity of the shorter sgRNAs, continued to accumulate insertions to an average length of 8.4 generated bps, encoding 15.3 bits of information (**Figure 4B-C**). In short, through DNA writing, the CHYRON_16i_ locus progressively generated far more information at a single site than any previous erasing-based DNA recorder and should be broadly useful.

**Figure 4.**
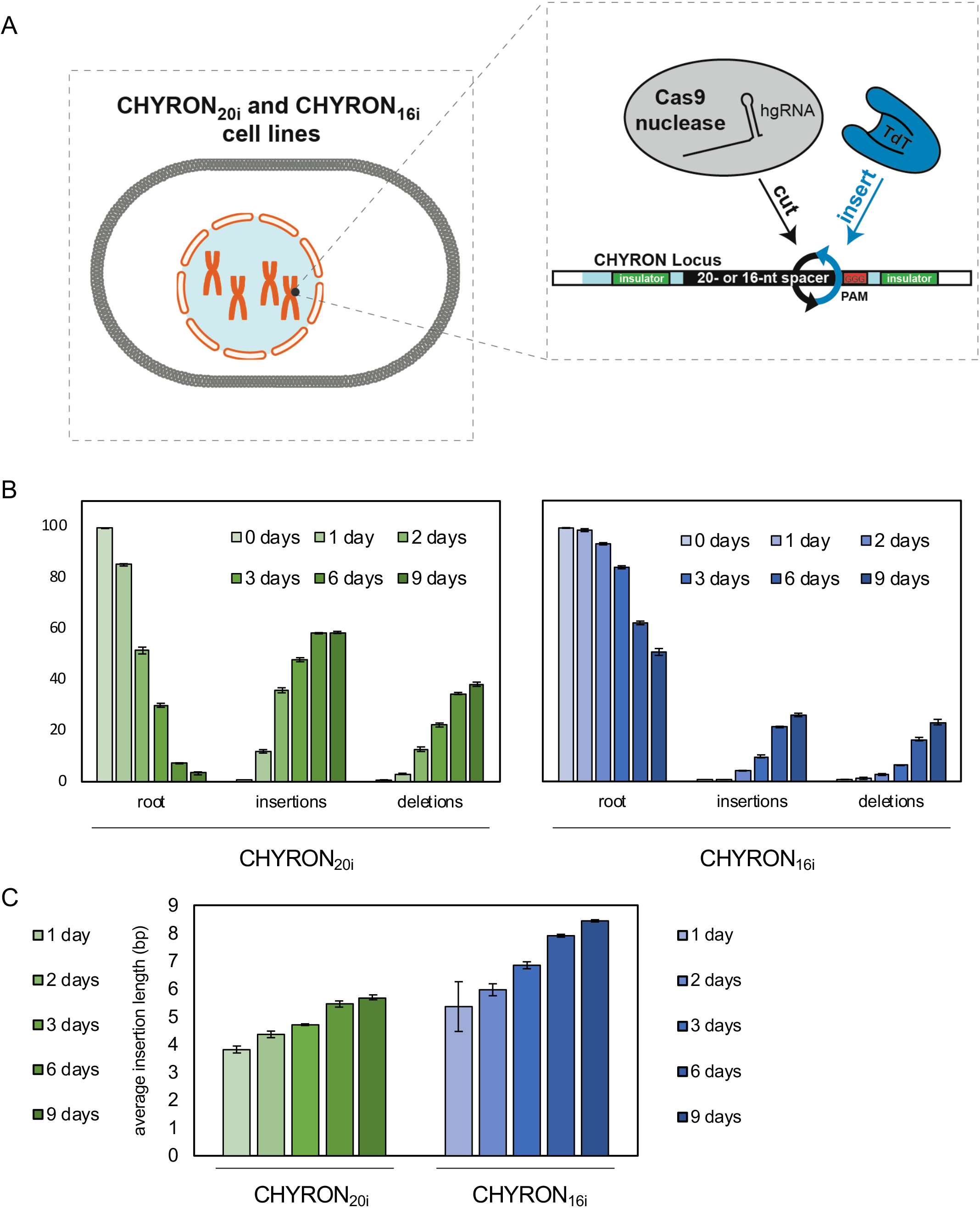
An improved version of CHYRON writes an average of 8.4 bps in an average of three rounds. **(A)** Clonal 293T cell lines (hereafter 293T-CHYRON_20i_ and 293T-CHYRON_16i_), were created by integrating cassettes at the *AAVS1* safe harbor locus. The cassettes contain hgRNAs with initial lengths of 20 or 16 nt, flanked by insulator sequences to prevent silencing. **(B)** CHYRON loci with an initial hgRNA length of 20 nt accumulated insertions and deletions over 6 days, whereas those with an initial hgRNA length of 16 nt accumulated insertions and deletions more slowly, continuing to do so through a 9-day timecourse. 293T-CHYRON_20i_ and 293T-CHYRON_16i_ cells were transfected with a plasmid expressing Cas9 and TdT for the indicated time before collection. Cells were re-transfected every 3 days. The CHYRON locus was analyzed by NGS and each sequence was annotated as root, pure insertion (insertions) or any sequence that involves a loss of information (deletions). Bars represent the average of three technical replicates. Error bars=±stdev. **(C)** Insertions continued to grow throughout the 9-day timecourse. The average lengths of pure insertions were calculated from the experiment in B. Bars represent the average of three technical replicates. Error bars=±stdev.

### CHYRON_16i_ allows the reconstruction of relationships among 16 populations containing thousands of lineages

To mimic a process of growth and spatial expansion over several days in a setting in which we could know the ground truth of lineage relationships among the cells, we (1) grew ~10,000 293T-CHYRON_16i_ cells (**Figure 5A**) bearing the root CHYRON_16i_ sequence in a well, (2) expressed Cas9 and TdT for three days (approximately three doublings) to allow the cells to write new bps at the CHYRON_16i_ locus, (3) split the well into two, and (4) repeated steps 2 and 3 again to yield 4 final wells (**Figure 5B**). We performed this process in quadruplicate to yield 16 final wells total. Cells in these final wells were allowed to grow for three days. The approximately three doublings between splits ensured that enough cells could acquire an insertion and then divide between splits, and the grow-out at the end of the experiment ensured that our recovery of unique sequences was high.

**Figure 5.**
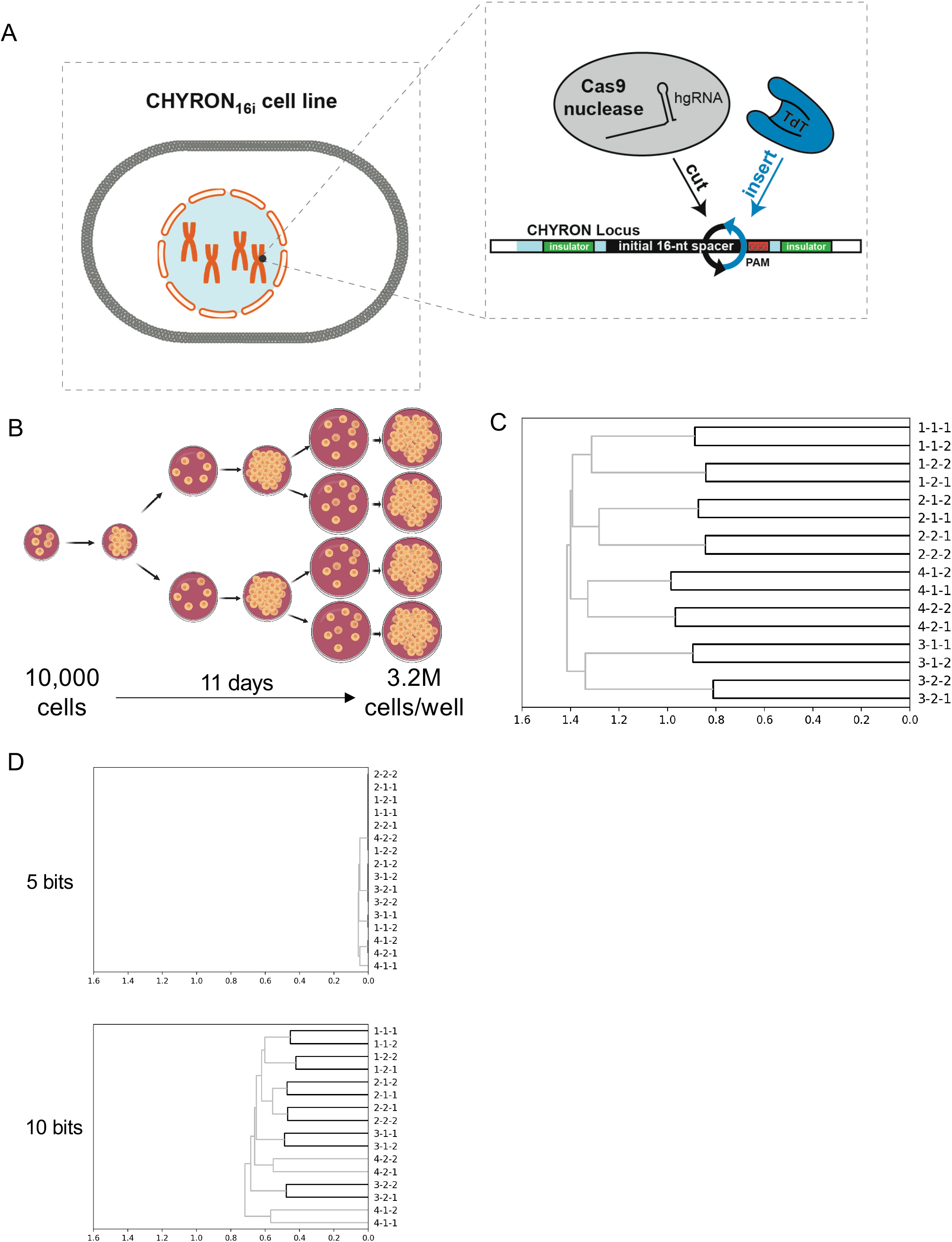
Reconstruction of cell relatedness by sequencing of the CHYRON locus. **(A)** The CHYRON_16i_ cell line was used. **(B)** Plan of the experiment. This experiment was performed in quadruplicate, such that 16 wells were collected and sequenced at the end of the experiment. Cells were transfected with Cas9 and TdT one day after each plating. **(C)** A simple method led to perfect reconstruction of the relatedness of all wells. For each well, a list was created of all unique insertions with an abundance of at least 0.0139% of the non-deletion reads and a length of 8-15 bp. Next, the total number of identical sequences between each pair of wells was divided by the total number of sequences in those wells that are not identical to compute the Jaccard similarity coefficient between that pair of wells. Then, hierarchical reconstruction was performed using the UPGMA algorithm. Units are arbitrary, but consistent across all plots in this work. **(D)** Reconstruction of these lineages would have been impossible with a recorder encoding 5-10 bits. The process in (C) was repeated after the data were transformed so that each locus encoded a maximum or either 5 or 10 bits.

By subjecting cells in only the final wells to NGS of the CHYRON locus, we were able to robustly generate a perfect reconstruction of the full splitting procedure. This was done using the presence of shared sequences between pairs of wells to calculate relatedness (similarity) and a standard agglomerative hierarchical clustering method to generate the tree from pairwise similarities (**Figure 5C**). The ease and accuracy of lineage reconstruction in this rather complex experiment – one population expanded to sixteen, and 10^4^ cells expanded to ~4 x 10^7^ (**Figure 5B**) – suggests that CHYRON should be a powerful system for deep lineage reconstruction of developmental processes involving substantial proliferation and fate changes.

It is instructive to note that perfect reconstruction resulted from the analysis of CHYRON insertions at least 8 bp in length (**Figure 5C**) and the quality of the reconstruction decreased slightly when including shorter insertion sequences *(e.g.* 7 bp, **Figure S6A**). In addition, the exclusion of shorter sequences was essential for accurate reconstruction when the abundance cutoff was set low (**Figure S6B**) or when the data were downsampled (**Figure S6C**). Why did the inclusion of shorter insertion sequences reduce reconstruction robustness? The answer to this question results from the interplay between convergence and sampling, an observation that should apply to all DNA-recording-based lineage reconstruction studies.

Let us consider a well (Well A), its closest relative (Well B), and a totally unrelated well (Well X). Suppose we detect a specific CHYRON sequence (or a shared substring of the CHYRON sequence given the ordered nature of insertions in CHYRON) in only two of these three wells. What is the chance that the two wells are Well A and Well B? If the sequence is long, then the chance is high, since a long CHYRON sequence is unlikely to arise independently. However, if the sequence is short, then it is possible that the two wells sharing the sequence are, for example, Well A and Well X, and that the sequence was generated independently in both wells. The critical issue, however, is that the sequence could also be absent in Well B, because (1) the sequence was generated after Well A and Well B split from their common ancestor or (2) the sequence was present also in Well B but was not detected due to sampling inefficiencies. In that case, the short sequence will preferentially assign relatedness to the unrelated wells over the related wells, reducing the accuracy of reconstruction. These considerations are treated further in **Supplementary Discussion**. We observe that potentially informative sequences, those that were generated before Well A and Well B split, and are successfully sampled from each well, decline with the square of the proportion sampled. However, potentially misleading sequences, that were generated independently in, for example, Well A and Well X, decline only linearly with decreased sampling. Therefore, one must use sufficiently long CHYRON sequences and ensure sufficient sampling of the sequences in all wells, a conclusion generalizable to all lineage reconstruction studies from self-mutating DNA recording systems. Longer, more information-dense sequences can partially, but not completely, compensate for poor sampling.

In order to compare CHYRON to previously-described DNA recording systems that record at a single site, we attempted to more exactly characterize the information used in the CHYRON reconstruction. We took advantage of the large and deeply-sequenced mock lineage tracing dataset to more precisely quantify the average entropy of insertions at the CHYRON locus (see **Supplementary Discussion** and **Table S6**). We found that the average entropy of CHYRON_16i_ in this experiment was 1.73 bits per bp. In this lineage reconstruction experiment, the average length of insertions observed was 7.8 bp (**Table S4**) so the expected value for the entropy per each generated CHYRON sequence was 13.5 bits. We tested whether a recorder that encoded less information could have enabled reconstruction of the relationships between the cell populations reconstructed in **Figure 5C**. To do so, we computationally reduced each insertion in the dataset to its first 3-bp (with an estimated information-encoding capacity of approximately 5 bits, the capacity of an hgRNA with Cas9 alone^4^) or 6-bp (approximately 10 bits) sequence. In neither case was a perfect reconstruction possible (**Figure 5D**). When insertions were limited to the first 3 bp, all 84 possible 1-, 2-, and 3-bp sequences were represented in each well (see github.com/liusynevolab/CHYRON-lineage for all data and analysis software). Thus, relationships between populations of these sizes cannot be reconstructed with a recorder encoding only 5 bits. All possible 6-bp sequences were not observed, but sequence convergence likely prevented a perfect lineage reconstruction for the 10-bit recorder as well. Successful lineage reconstructions with lower-information DNA recorders have used either smaller populations^4,12^ or multiple recording sites^2,5,6,12,13^.

### CHYRON_20_ and CHYRON_16_ can report the dose and duration of exposure to a hypoxia mimic

DNA recording systems have been used to log cellular exposure to biological stimuli by making mutation at the recording locus inducible by biological stimuli of interest. For example, mutational accumulation at hgRNAs have been linked to inflammation exposure^3^. However, in such cases, recording has been digital from the perspective of a single cell: the information used from the hgRNA is whether it is mutated or not. The dose or duration of the stimulus being recorded is therefore reflected at the population level, in the proportion of the population that bears a mutated hgRNA. While CHYRON can do the same, CHYRON also offers the possibility of acting as a compact recorder that is analog from the perspective of a single cell. This is because the CHYRON locus progressively writes new bps and rarely erases, so a CHYRON locus should grow monotonically longer as the cell is exposed to the stimulus for a longer period of time, or at a higher dose.

To test stimulus recording with CHYRON, we linked insertional mutagenesis at a CHYRON locus to hypoxia, which triggers adaptive responses that affect a wide range of cellular behaviors, including several, such as migration and invasion, that are important for tumor evolution and metastasis^28,29^. We created a construct in which the expression of Cas9 and TdT is under the control of the 4xHRE-YB-TATA^30^ promoter, and in which Cas9 is additionally fused to an oxygen-dependent degron domain^29^ (**Figure 6A**). We transfected 293T-CHYRON_20_ and 293T-CHYRON_16_ (which bears a 16-nt-spacer hgRNA integrated at the *AAVS1* locus, as in CHYRON_16i_ but without insulators) with this construct, and then exposed them to three different concentrations of the hypoxia mimic DMOG for five different durations. In both cell lines, the proportion of the population bearing insertions increased with dose and duration of DMOG treatment (**Figures 6B**). In the case of CHYRON_16_, the average length of insertions also increased with duration of DMOG exposure for the sequences that were mutated (**Figure 6C**). In other words, CHYRON is capable of recording exposure to stimuli in a manner that is digital (**Figures 6B**) or analog (**Figure 6C**), where the latter of these modes can in principle provide information on the experience of each single cell. We note that currently, the dynamic range of analog recording achieved with CHYRON is narrow (**Figure 6C**), but with further development, we expect CHYRON will be an ideal system for capturing detailed cellular histories at single-cell resolution.

**Figure 6.**
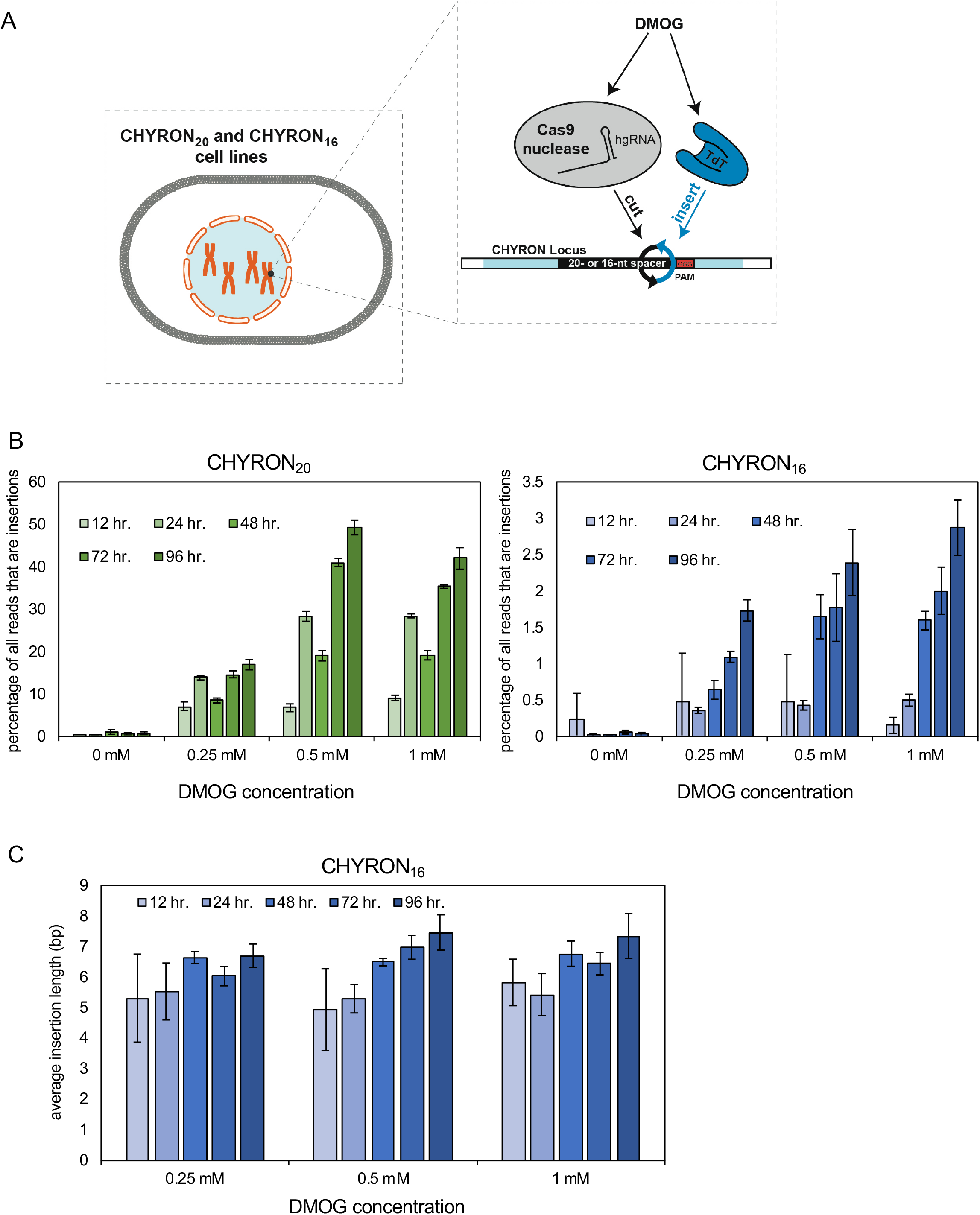
Expression of hypoxia-inducible Cas9 and TdT promotes insertion abundance and length in proportion to duration of treatment with the hypoxia mimic DMOG. **(A)** These experiments were performed using the CHYRON_20_ and CHYRON_16_ cell lines. **(B)** When transfected with hypoxia-inducible Cas9 and TdT, an ever-higher proportion of CHYRON_20_ loci (left) accumulated insertions upon longer treatment and higher doses of DMOG, a hypoxia mimic, whereas CHYRON_16_ loci (right) accumulated insertions at a lower, but still dose-dependent, rate. 293T-CHYRON cells were transfected with a plasmid encoding Cas9 and TdT under the control of a promoter containing four copies of the hypoxia response element; Cas9 is additionally fused to a degron that destabilizes proteins in the presence of normal levels of oxygen. After transfection, cells were treated with DMOG or a vehicle control and then collected at the indicated time and analyzed as in Figure 3B. Bars represent the average of three technical replicates. Error bars= ± stdev. **(C)** In 293T-CHYRON_16_ cells transfected with hypoxia-inducible Cas9 and TdT, insertions grew longer with increasing duration of exposure to DMOG. Lengths of pure insertions in the CHYRON_16_ experiment were calculated. Bars represent the average of three technical replicates. Error bars=±stdev.

### Current and future capabilities of CHYRON

There are three unique features of the CHYRON architecture that we believe will lead to its broad application and motivate its continued development in our and other labs. First is the high information content and density of CHYRON. CHYRON is able to diversify a very compact recording locus, consisting of a single site that is repeatedly modified, so that the locus can bear a unique sequence in each of tens of thousands of cells. This capability may be especially important for applications where it is difficult to capture all cells that might be related to each other, in which case a DNA recorder with a high information content is necessary to limit the possibility of misleading convergent sequences in unrelated cells. Second is the property that CHYRON records information by generating an ordered accumulation of random insertions. Unlike deletions and substitutions, pure ordered insertions gain information without corrupting or removing previous information, which is ideal for a DNA recorder. The ordered nature of insertions generated also introduces the possibility that, if TdT can be engineered to add different types and lengths of nts deterministically and the activity of the different TdTs can be coupled to different cellular stresses or the cell cycle, the CHYRON locus would record the relative timings of the different stresses in the cell’s history or even provide an accurate count of cell divisions. The latter may enable single-cell-resolution lineage reconstruction from sparsely-sampled CHYRON sequences, a goal we are actively pursuing. Third, CHYRON is a high-information DNA recorder that uses only a single genomic site. Because the recording site and Cas9/TdT machinery can be encoded at a single locus, CHYRON can be readily transplanted into new cell or animal lines.

In its current form, CHYRON is likely the best option for reconstruction of the relationships between thousands of cell lineages resulting in millions of cells. However, additional work is necessary to reach its full potential. Currently, TdT is recruited to the Cas9 cut site through its natural interaction with the DSB repair machinery. As a result, it is likely recruited to all DSBs in the cell and will act as a mutagen over long periods of expression. To ensure that normal development is not perturbed by this increased mutagenesis, TdT should be engineered so that it is specifically recruited only to the CHYRON locus. As shown in this study, we could not observe increased activity of TdT *in cis* when it is fused to Cas9 (**Figure S2C**). Mutations of TdT that prevent its binding to the DNA repair machinery have been reported^31^, and a better strategy may be to fuse these mutants to a protein that remains at the DSB after Cas9 dissociates. The information-encoding capacity of CHYRON, although unprecedentedly high for a single site, is limited by the declining efficiency of the hgRNA as it grows longer. The reduced efficiency likely arises from a combination of guide RNA length and secondary structure in the critical seed region. Several approaches could address these issues, namely engineering Cas9 to better tolerate these types of sequences or using a different nuclease that cuts further from its seed region. Finally, the ~25% rate of deletion per Cas9 cut will still eventually lead to information loss and inactivation of all CHYRON loci in the limit of truly continuous recording. However, recruitment of factors that manipulate the balance of DSB repair pathways at the Cas9 cut site could reduce deletions significantly. The future development of CHYRON will be enhanced by the wide interest in engineering new capabilities into its protein components – a CRISPR nuclease^8^ and TdT, in which there has been considerable recent interest as a tool for *in vitro* DNA synthesis^32,33^. Techniques that use polymerases^34,35^, including TdT^36^, to record time-series information on DNA synthesis timescales *in vitro* could also be merged with CHYRON. In short, we predict that the uniqueness of CHYRON as a DNA recorder based on writing DNA rather than erasing DNA and the promise of CHYRON in reaching fully continuous recording of biological stimuli or lineage relationships at single-cell resolution *in vivo* will spur its continued development and application.

## Materials and Methods

### Plasmid cloning

Cloning was done by standard Gibson assembly, *in vivo* recombination, and restriction-ligation cloning. All plasmids are listed in **Table S7**, and available along with full sequences at Addgene (addgene.org/browse/article/28203329). Detailed descriptions of cloning procedures can be found in **Supplementary Methods**. All plasmids to be used for transfection were purified with HP GenElute Midi or Mini kits (Sigma # NA0200 and NA0150).

### Cell culture and transfection

All cell culture experiments were performed in HEK293T cells obtained from ATCC (CRL-3216). Cells were cultured in DMEM, high glucose, GlutaMAX™ Supplement (Gibco #10566024), supplemented with 10% FBS (Sigma #12306C), at 37°C and 5% CO_2_.

Transient transfections were performed by mixing DNA with Fugene (Promega #E2311) in serum-free DMEM, at a ratio of 1 μg DNA to 3 μL Fugene

To create the 293T-CHYRON cell lines, the plasmid to integrate the hgRNA into 293T cells was digested with EcoRI-HF (NEB#R3101), then purified on a silica column (Epoch #3010). 350 ng of this plasmid was mixed with 100 ng of MSP680, a plasmid expressing Cas9^EQR^, a gift from Keith Joung (Addgene #65772)^44^, 50 ng of a plasmid expressing an sgRNA against a sequence at HEK293 site 3 or *AAVS1* that can be cut by Cas9^EQR^, and 1.5 uL Fugene. 293T cells were transfected in a 24-well dish, transformants were selected with 1-2 ug/mL of puromycin (Invivogen #ant-pr-1), then a single colony was isolated in two rounds of dilution and colony picking.

Samples of all cell lines used in this study, including 293T and 293T-CHYRON cells, corresponding to the latest frozen stock that was used, were commercially tested for mycoplasma contamination and shown to be negative (Applied Biological Materials, Inc.).

### Examining the insertion bias of TdT at varying cut sites

To test the insertion characteristics of TdT, 16 targetable sites were chosen that contained all combinations of each nucleotide at the −4/-3 position relative to the PAM (Supplementary Table 1). A six-well well of HEK293T cells were transfected with 0.4 μg of the specific sgRNA and either 0.92 μg of Cas9, 1.09 μg of Cas9-T2A-TdT, or 1.1 μg of Cas9-5XFlag-TdT. To normalize the total amount of DNA transfected, the Cas9 and Cas9-T2A-TdT transfections were supplemented with 0.12 μg and 0.01 μg of pcDNA3.1-sfGFP, respectively. The cells were collected three days post-transfection and processed for DNA and protein as detailed below.

### Long-term editing on the CHYRON locus

293T-CHYRON_20, 20i_, or _16i_ cells were transfected in 6-well dishes with 2 μg of the Cas9-T2A-TdT construct. The editing took place for 1, 2, 3, 6, or 9 days after transfection. For the 6 and 9-day time point, 10% of the cells from the previous time point were used to seed a new culture to be transfected again with the same amount of DNA as the previous transfection. As a control, the same experiment was performed with a Cas9-T2A-TdT plasmid with a stop codon five amino acids into the TdT sequence. All time points were collected as a single well of a six-well plate (Falcon # 08-772-1B).

### Two-step editing with Cas9 and TdT via isolation of single colonies

293T-CHYRON_20_ cells were transfected with equal amounts of plasmids expressing Cas9-5xFlag-TdT and free TdT, then diluted and single colonies picked. The CHYRON locus of these colonies was sequenced by the Sanger method, and six cell lines were chosen for further study. These six cell lines were grown in two 6-wells each. For each, one well was transfected with a plasmid expressing Cas9-T2A-TdT and the other well was untransfected. Three cell lines representing two insertions were found to be nearly clonal and successfully sequenced. All samples were collected and the CHYRON locus sequenced via UMI incorporation and NGS.

### Lineage reconstruction assay and analysis

5,000 293T-CHYRON_16i_ cells were plated in each of 4 wells of a 384-well plate, then transfected the next day with a plasmid expressing Cas9 and TdT (pcDNA-Cas9-T2A-TdT). Three days later, when they had expanded to approximately 86,000 cells per well, they were each split into two wells of a 24-well, allowed to attach for one day, then transfected again. Three days later, when each well had expanded to approximately 800,000 cells, each well was split into two wells of a 6-well dish. Three days later, all wells were collected and analyzed by amplicon sequencing without UMI incorporation.

For our initial analysis, we created a list of all insertion sequences in each well. Each insertion has an “abundance,” based on the number of NGS reads that include that exact insertion sequence, and a length, equal to the number of bp added to the root sequence at the Cas9 cut site. We refined the list for each well to include only those insertions that met two criteria: (1) they were represented in at least 0. 0139% of the non-deletion reads in the well and (2) the inserted sequence had a length of 8 to 15 bp. From this list of insertions, we created a binary vector for each well whose length was equal to the total number of insertion sequences with these criteria observed in any of the 16 wells in the experiment. The vector for each well contains a 1 for a particular insertion if that insertion was present in the refined list for that well, or a 0 if that insertion is absent. We used these vectors to calculate the Jaccard similarity between each pair of wells^4^, then reconstructed the relationships using the UPGMA hierarchical clustering algorithm (github.com/scipy/scipy/blob/v1.2.1/scipy/cluster/hierarchy.py#L411-L490).

All the analyses were done in Python. The scripts are available at github.com/liusynevolab/CHYRON-lineage.

### Hypoxia recording assay

293T-CHYRON_20_ and 293T-CHYRON_16_ cells were transfected in 6-well dishes with 2 μg of the 4XHRE-YBTATA-Cas9-ODD-T2A-TdT construct. Ten hours after transfection, fresh medium supplemented with 0, 0.25, 0.5, or 1 mM DMOG (EMD Millipore Calbiochem™ #40-009) was added. Cells were collected and DNA extracted at 24 or 48 hr. after DMOG addition. At 48 hr., cells were replated and retransfected 14 hours later. 14 hours after transfection, DMOG was added at the indicated concentrations, then the cells were grown for 24 hr. before collection of the 72 hr. timepoint, and 48 hr. before collection of the 96 hr. timepoint.

### Deep sequencing library preparation of a genomic locus (for Figures 2, S1, and S2)

Genomic DNA was isolated with a QIAAmp DNA Mini Kit (Qiagen #51304) and the region targeted by Cas9 was amplified by PCR. The primers contained the Illumina adapters and a 5 – 7 nt sample-specific barcode (Supplementary Table 2). The PCR reaction was performed with Q5 Hot Start High-Fidelity DNA Polymerase (NEB) and the following protocol: 98°C, 1 min; (98°C, 10 s; 60°C, 30 s; 72°C, 30 s) × 35; 72 °C, 1 min. Each reaction was done with 100 ng of nucleic acid. For the same genomic locus, each sample was normalized by signal intensity on a 0.9% gel and pooled into a single mixture, which was cleaned using a NucleoSpin Gel and PCR Clean-up Kit (Macherey-Nagel #NC0389463). 10 ng per individual sample from the pooled clean product was sent to Quintara Biosciences or FornaxBio and ran on an Illumina MiSeq.

At the sequencing vendor, the libraries were purified by binding to AMPure beads (0.9 beads:1 sample) and further amplified to incorporate the TruSeq HT i5 and i7 adaptors, using Q5 High Fidelity DNA Polymerase, for 10-13 cycles. The amplified libraries were agarose gel-purified, including at least 100 bp of room around the desired bands, to avoid biasing against deletions or insertions, and then sequenced on an Illumina MiSeq using the 500-cycle v2 reagent kit (Illumina Cat # MS-102-2003).

### Deep sequencing library preparation of the CHYRON locus at high efficiency for lineage reconstruction (for Figures 5, S4, S5, S6, and S7)

Genomic DNA was extracted with the QIAmp DNA Micro Kit (Qiagen) and carrier RNA (for Figure S4) or the QIAmp DNA Mini kit and the entire recovery was used in the initial PCR. The initial PCR was performed for 25 cycles with Phusion HotStart Flex polymerase in GC buffer (New England Biolabs), each sample was purified with AMPure beads (0.9:1), then digested with PmlI (New England Biolabs) for 4 hours, then purified with AMPure beads again. Then the reamplification PCR was performed for 15-25 cycles in Q5 HotStart polymerase. The samples were pooled according to their estimated concentration on an agarose gel stained with ethidium bromide, then purified with AMPure beads, and cut with PmlI for an additional 4 hours. Finally, bands of the expected library size or up to 100 bp larger were gel-purified using a Macherey-Nagel PCR Clean-up kit. They were sequenced on an Illumina HiSeq 2500 using the PE100 kit for Figures 5, S5, S6, and S7, or on an Illumina MiSeq using the PE250 kit for Figure S4, at the UCI Genomics High Throughput Facility.

### Unique molecular identifier incorporation for sequencing of the CHYRON locus (for Figures 3, 4, 6, and S3)

To barcode individual cells containing the integrated CHYRON locus, unique molecular identifiers (UMIs) of 20 degenerate nucleotides were incorporated. Primers were ordered from IDT containing the following: the Illumina reverse adapter, a UMI, a 5 – 7 nt sample-specific barcode, and a stgRNA construct binding region (**Table S3**). gDNA was isolated with the QIAmp DNA Mini Kit (Qiagen) and 600 ng of nucleic acid was used. The UMI incorporation reaction was run with Phusion Hot Start Flex DNA Polymerase (NEB) and under the following condition: 98°C, 5 min; (55°C, 30 s at a ramp rate of 4°C/s ramp rate; 72°C, 1.5 min) × 10. The reaction was enzymatically cleaned with Exonuclease I and Shrimp Alkaline Phosphatase (NEB) by incubating the sample and enzymes for 30 minutes at 37°C.

A downstream PCR was performed on the UMI incorporation step to amplify specific sequences that contained a UMI. The sample was run with a forward primer with the Illumina forward adapter, a 5 – 7 nt sample-specific barcode, and a stgRNA binding region, and a reverse primer complementary to the Illumina reverse adapter present on the UMI primer. The PCR was performed under the following conditions: 98°C, 3 min; (98°C, 1 min; 65°C, 30 s; 72°C, 30 s) × 35; 72°C, 1 min with a 2°C/s ramp rate. Products were purified on columns from the NucleoSpin Gel and PCR Clean-up Kit (Macherey-Nagel) and individual samples were pooled based on equal molar ratios. Samples were further processed as for genomic sites for **Figure 3A-C**. For the rest of the experiments, samples were individually purified with AMPure beads (0.9:1), then reamplified for 15 cycles, pooled, and gel-purified including ~50 bp smaller and 100 bp larger than the expected band. Libraries were sequenced at the UCI Genomics High-throughput Facility on an Illumina MiSeq using the 500-cycle v2 reagent kit (Illumina Cat # MS-102-2003).

### Deep sequencing analysis

The sequences retrieved by next generation sequencing were first grouped to individual samples based on their barcodes. Then for each sample, associated forward and reverse reads were merged (Pear 0.9.10^46^) and mapped to the reference sequence by the alignment algorithm implementation (Mapp) used in Perli *et al.,* 2016^3^ which provides a sequence of M(Match), X(Mismatch), I(Insertion) and D(Deletion) as the mapping result.

If UMI barcodes are present, before mapping, sequences with the same barcodes are combined to one. To combine, we started with a multiple-alignment of the sequences (done by Motility library in Python) with the same UMI barcode (One nucleotide difference was allowed in UMI barcodes) and to avoid any random mismatches produced in the sequencing process, for each position in this alignment only nucleotides present in more than 50% of the sequences in this group, were used to generate the consensus sequence.

After the alignment, first bad alignments (>20 mismatches or >50% deletions) were removed. Then, mismatches and inserted or deleted sequences, their positions on the reference sequence and their frequencies were extracted. Only insertions or deletions occurring around (−10nt to +10nt) the cut side were kept. (For the data in Figures 2 and S2, we used the region −7 bp to +7 bp from the cut site to avoid a genomic SNP.) To remove insertions that were the result of homologous recombination repair, if longer insertions(>12nt) could map (with less than 2nt difference) to sequences of the plasmids that were transfected, they were filtered out. In addition, insertions longer than 15 nt (or 20 nt for Figure 2C and 40 nt for S4) were excluded, as these were found to more frequently have nucleotide biases that suggested they were TdT-independent (data not shown).

All the analyses were done in Python. The scripts are available at github.com/liusynevolab/CHYRON-NGS.

For the experiment shown in Figures 5, S5, S6, and S7, because they were sequenced with a paired-end 100 protocol, rather than the paired-end 250 we used for all other experiments, forward and reverse reads were not paired, and forward reads only were used for the analysis.

### Figure preparation

Figures 5B, S4A, and S4E were prepared at biorender.com. The plots in Figures 5C, 5D, S4B, S4F, and S6A were generated using the hierarchy.dendrogram function in matplotlib (scipy.org). The threshold for color change was set at the default level: 0.7*max(Z[:,2]).

### Availability of data and reagents

All NGS data sets have been deposited at the NCBI’s Sequence Read Archive, accession # PRJNA561027. Plasmids have been deposited at Addgene. See Table S7 for a guide to these reagents. Please contact CCL for cell lines.

### Code availability

All scripts are available at github.com/liusynevolab.

## Abbreviations

CRISPR: clustered regularly interspaced short palindromic repeat,
DNA: deoxyribonucleic acid,
CHYRON: cell history recording by ordered insertion,
bp: base pair (of DNA),
TdT: terminal deoxynucleotidyl transferase,
hgRNA: homing guide ribonucleic acid,
stgRNA: self-targeting guide RNA,
Cas9: CRISPR-associated 9,
GFP: green fluorescent protein,
DSB: double-strand break (in DNA),
nt: nucleotide (of DNA or RNA),
sgRNA: single-guide RNA,
stdev: standard deviation.

## Acknowledgements

We thank Seanjeet K. Paul and Tate C. Lone for technical assistance. We thank the following people for helpful discussions: Christian Guerrero-Juarez, Chengjian Li, Jan Zimak, Qing Nie, and all members of the Liu Laboratory. We thank the following people for plasmids: Yvonne Chen, Keith Joung, William Kaelin, George Church, David Liu, Eric Campeau, Paul Kaufman, and Timothy Lu. This work was made possible, in part, through access to the Genomics High Throughput Facility Shared Resource of the Cancer Center Support Grant (P30CA-062203) at the University of California, Irvine and NIH shared instrumentation grants 1S10RR025496-01, 1S10OD010794-01, and 1S10OD021718-01. This work was funded by NIH grants 1DP2GM119163-01 and 1R21GM126287-01 to CCL, an AHA Predoctoral Fellowship to CKC, and a fellowship from the NSF-Simons Center for Multiscale Cell Fate Research (NSF Award #1763272) to TBL.

## Author contributions

TBL and CCL designed experiments. TBL, JHG, MWS, and CKC performed experiments. TBL, JHG, GL, and CCL developed protocols. TBL, JHG, and CKC made cell lines. TBL, JHG, CKC, and GL made and validated reagents. TBL, MWS, EF, BSA, XX, and CCL designed and performed analyses. MWS, EF, BSA, BL, MF, and TBL wrote code. TBL and CCL wrote the paper, with input from all authors, especially CKC. CCL procured funding and oversaw the project.

## Supplementary Material Contents

Supplementary Methods

Supplementary Tables

Table S1. All nucleotide bias results underlying Figure 2C, including target sites and NGS primers, and Shannon entropy calculations.

Table S2. Data underlying Figure S4A-D.

Table S3. Data underlying Figure S4E-F.

Table S4. Data underlying Figures 5, S5, S6, and S7.

Table S5. Expanded version of Figure S7.

Table S6. Tabulation of all 1- and 2-nt sequences in the data underlying Figures 5, S5, S6, and S7, and Shannon entropy calculations based on these results.

Table S7. Guide to plasmids available at Addgene, NGS datasets available at the NCBI Sequence Read Archive, and NGS primers and reference sequences.

## Supplementary Discussion

Supplementary Figures

Figure S1. Expression of Cas9 upon transfection, effect of TdT on editing outcomes at a variety of genomic sites, and test of fidelity of NGS library prep for GC-rich templates. Related to Figure 2 and Methods.

Figure S2. Activity of varying amounts and fusions of TdT. Related to Figure 2.

Figure S3. Further characterization of CHYRON_20_. Related to Figure 3.

Figure S4. Robust lineage reconstruction with CHYRON_20_ is only possible for splitting protocols with good sampling efficiency. Related to Figure 5.

Figure S5. Lineage reconstruction using CHYRON-_16i_ is robust. Related to Figure 5.

Figure S6. High-information recording and optimal analysis is required when sampling is limited. Related to Figure 5.

Figure S7. Analysis by alphabetizing of insertions at the CHYRON-_16i_ locus. Related to Figure 5.

