## Supplementary material for "DNA writing at a single genomic site enables lineage tracing and analog recording in mammalian cells": Supplemntary Information

### Supplementary Methods

**Plasmid cloning.** Polymerase Chain Reactions (PCRs) were performed with Q5 Hot Start High-Fidelity DNA Polymerase or Phusion Hot Start Flex DNA Polymerase (New England BioLabs). All primers were purchased from Integrated DNA Technologies (IDT) and PCR reagents were provided by NEB. All transformations were done in Top10 *E. coli* (ThermoFisher Scientific), unless otherwise stated.

The human codon-optimized *S. pyogenes* Cas9 DNA sequence was PCR amplified from hCas9, which was a gift from George Church (Addgene plasmid # 41815[21]). Either an XTEN linker or a T2A self-cleaving sequence along with TdT were cloned onto the C-terminus of Cas9. The XTEN linker was cloned through PCR from pCMV-BE3, which was a gift from David Liu (Addgene plasmid # 73021[37]). The T2A self-cleaving sequence was inserted through PCR by designing primers with overhangs containing the T2A sequence. The TdT coding sequence was amplified from the cDNA of an acute lymphoblastic leukemia cell line through PCR and this entire insert was cloned into a pcDNA3.1 backbone, yielding Cas9-XTEN-TdT or Cas9-T2A-TdT. Cas9-containing constructs with the pcDNA3.1 backbone were transformed into XL10-Gold Ultracompetent *E. coli* (Agilent #200315).

Additional Cas9-TdT constructs containing different linkers were cloned through restriction enzyme digestion and ligation. All restriction enzymes and T4 DNA Ligase were purchased from NEB. All vector digestions were treated with alkaline phosphatase, calf intestinal (CIP) from NEB after complete digestion by the restriction enzymes. A base construct was cloned first through Gibson Assembly to add restriction sites to the original Cas9-XTEN-TdT construct: <u>NheI</u>-<u>SfiI</u>-Cas9-<u>KpnI</u>-XTEN-<u>SexAI</u>-TdT. The base construct was digested with KpnI and SexAI. Three separate PCR's were performed to yield a 5xFlag or 5xGSA linker product with KpnI and SexAI restriction sites. The 5xFlag was present on a gBlock (IDT) and the 5xGSA linker (4 repeats of the sequence GSAGSAAGSGEF and a final repeat with the sequence GSAGSAAGASGEGRP[38]) was ordered on a minigene (IDT). These two inserts were digested with the appropriate enzymes and ligated individually into the KpnI and SexAI digested base construct yielding Cas9-5xFlag-TdT or Cas9-5xGSA-TdT.

Additional PCR on the 5xFlag gBlock yielded 5xFlag flanked by KpnI sites, which was digested and then ligated into Cas9-5xFlag-TdT, resulting in Cas9-10xFlag-TdT. To clone Cas9-15xFlag-TdT, a subsequent PCR was performed on the gBlock to amplify the 5xFlag sequence with FseI restriction sites. The FseI restriction-enzyme recognition sites were used to ligate the 5xFlag sequence into Cas9-10xFlag-TdT, yielding Cas9-15xFlag-TdT.

Hypoxia inducible constructs were cloned through adding a 4x hypoxia-response element (HRE)-YBTATA promoter to drive the expression of Cas9-T2A-TdT. The 4xHRE sequence was PCR amplified from 4xHRE_v2_YB-TATA-Gluc-CMV_dsRed[30], which was a gift from Yvonne Chen. In addition, the oxygen-dependent degron (ODD) was cloned onto the C-terminus of Cas9. The ODD sequence was amplified from HA-HIF1alpha-wt-pBabe-puro, which was a gift from William Kaelin (Addgene plasmid # 19365[39]). This plasmid also includes blasticidin resistance (not used in this work), which was cloned from pLenti CMV Blast DEST (706-1) backbone, which was a gift from Eric Campeau and Paul Kaufman (Addgene plasmid # 17451[40]).

Control plasmids were cloned containing catalytically-dead TdT (dTdT). The dTdT DNA fragment was prepared through introduction of the D343E and D345E mutations[41,42] into the wild-type TdT sequence. Two control plasmids were cloned through Gibson Assembly into the pcDNA3.1 backbone: Cas9-XTEN-dTdT and dTdT alone.

To clone single-guide RNA (sgRNA) plasmids, the spacer region of the desired sgRNA was inserted into the pSQT1313 expression plasmid, under the control of the U6 promoter, which was a gift from Keith Joung (Addgene plasmid # 53370)[43]. The desired spacer region was introduced by PCR, Gibson assembly, and subsequent transformation. Alternatively, a single PCR was performed on the parent plasmid to create a linear product with homologous ends. This linear piece was transformed into SS320 *E. coli* (Lucigen) to allow for recombination to yield the desired variant sgRNA plasmid.

The homing gRNA (hgRNA) constructs contained the HEK293 site 3 sgRNA cassette with a GGG (instead of GTT) at the 3'-end of the spacer region and the complementary mutations in the opposite site of the hairpin[3,4]. This sequence was amplified from the sgRNA plasmid with the corresponding spacer and the PAM-introducing mutations were present on the PCR primer. The resulting U6 promoter-hgRNA variant was cloned into a pcDNA3.1 backbone with the CMV promoter driving a puromycin-resistance gene. In addition, 750 bp regions homologous to the HEK293 site3 locus (for $CHYRON_{20}$) or *AAVS1* (for $CHYRON_{16}$, $CHYRON_{20i}$, or $CHYRON_{16i}$) was cloned upstream and downstream of the hgRNA and selection marker region of the plasmid. The sequences of the flanks were PCR amplified from HEK293T genomic DNA and an EcoRI restriction site was added on the 5'-end of the upstream flank and on the 3'-end of the downstream flank. These restriction sites allowed for linearization of the plasmid to be stably integrated into HEK293T cells upon transfection.

The pcDNA3.1-sfGFP construct was cloned through Gibson Assembly of the superfolder GFP gene from the yeast toolkit[45] into pcDNA3.1.

**Western blot.** For determining protein expression of our Cas9 constructs, western blots were performed by first lysing cell pellets with 1X RIPA buffer and a protease inhibitor cocktail (Roche #4693159001). After 30 minutes on ice, the lysis reaction was spun down and the supernatant was used in a BCA Reagent Assay (ThermoFisher Scientific #PI23225) to normalize for protein concentration. Upon normalization, the necessary volume of supernatant was added to LDS Sample Buffer (ThermoFisher Scientific #NP0007) with 0.2% βME (Fisher Scientific #BP176100).

All protein gels were 4-12% Bis-Tris Gels, 1.0 mm, 15-well (Invitrogen, Thermo Scientific #NP0323) and electrophoresis was performed in an XCell SureLock Mini-Cell Electrophoresis System (ThermoFisher Scientific #EI0001) with 1X MOPS Running Buffer according to the NuPAGE MOPS SDS Running Buffer recipe. Protein transfers were performed in a Mini Trans-Blot Cell (Bio-Rad #1703930) with a transfer buffer made according to the Bjerrum Schafer-Nielsen Buffer with SDS (Bio-Rad). Protein membranes and blotting paper were components of the EMD Millipore Blotting Sandwich Immobilin-P (Millipore Sigma #IPSN07852).

After transfer, the protein membrane was cut to create separate sections for Cas9 and actin blotting. An additional protein gel and transfer were performed for experiments that blotted for TdT. The respective membrane was incubated in either Guide-it Cas9 Polyclonal Antibody (Clontech #632607), α-TdT Antibody (Abcam #ab14772), or α-Actin Antibody (Abcam #ab14128) for 3 hours at room temperature. Western blots were then incubated with horseradish peroxidase-fused secondary antibodies. α-Rabbit IgG (Sigma-Aldrich #A0545) was used to bind the primary antibody of Cas9 and TdT, while α-Mouse IgG (R&D Systems #HAF007) was used to bind the primary antibody of actin. The western blot membrane was treated with Clarity ECL Western Blotting Substrate (Bio-Rad #1705061). Blots were scanned on a ChemiDoc Touch Imaging System (Bio-Rad #1708370).

**Alternative DNA extraction from mammalian cell culture.** An alternative protocol was developed for the extraction of genomic DNA from mammalian cells, and was used for experiments shown in Figure S2. A cell pellet was lysed in a lysis buffer consisting of 20 mM EDTA, 10 mM Tris pH 8.0, 200 mM NaCl, 0.2% TritonX-100, 200 μg/μL proteinase K. The lysis reaction was incubated at 65 °C for 10 minutes and a 1:4 mixture of 7X lysis buffer (Zymo #D4036-1) to water was added and the reaction was further incubated at 65 °C. The lysis reaction was neutralized with neutralization buffer (Zymo #D4036-2) and cell debris was spun out. EconoSpin columns (Epoch #1910) were used to capture the DNA from the supernatant and a wash step with PE buffer was included before elution of the pure genomic DNA in water.

**Amplification bias assay of varying polymerases.** Since the bias for TdT-mediated insertions is for the nucleotides G or C, it was important to test which polymerase would be optimal for amplifying GC stretches inserted on the hgRNA. Three gBlock Gene Fragments (IDT) were ordered with an insertion of 40 nucleotides at the -3 position of the hgRNA. The insertion was either 50%, 65%, or 80% GC rich. The amplification test was either performed with Q5 Hot Start High-Fidelity DNA Polymerase or Phusion Hot Start Flex DNA Polymerase (NEB). Reactions were performed with 10 ng of the individual fragments as well as with a 1:1:1 mix of all three fragments. Both forward and reverse primers contained a 5 – 7 nt sample-specific barcode and the appropriate Illumina adapters. The eight PCR's were performed with the following protocol: 98 °C, 3 min; (98 °C, 1 min; 55 °C, 30 s; 72 °C, 30 s) × 35; 72 °C, 1 min with a 2 °C/s ramp rate. Each PCR was normalized using agarose gel electrophoresis and equal amounts of each sample were pooled based on signal intensity. Final pools were cleaned on a NucleoSpin Gel and PCR Clean-up Kit column (Macherey-Nagel) and sent to Quintara Biosciences or FornaxBio and sequenced as above.

**Pulsing of Cas9 and TdT expression for Figure S4E-F.** Each well in a 96-well plate was transfected with 50 ng of the plasmid (pTBL716) expressing Cas9-ODD-T2A-TdT under the control of the 4xHRE-YBTATA promoter (with 0.15 uL Fugene per the manufacturer's instructions). 6 hours later, the promoter was activated by changing to medium with 1 mM DMOG. 24 hours after addition of DMOG, the DMOG-containing medium was removed and replaced with fresh medium.

**Lineage reconstruction.** To determine the average prefix between two wells, we assess each insertion in the first well, pair it with its closest match in the second well, and record the length of the initial identical sequence (prefix) shared by those two insertions. After we perform this process for each insertion in both wells, we average the prefix lengths to determine the average prefix between those two wells. After calculating the average prefix shared by each pair of wells, we pair the two wells with the longest average prefix, then those with the next-longest average prefix, etc, until all sibling wells are paired. Then, the identical sequences from each pair of wells are collected the create a "pooled well," and those pooled wells are paired by the same process to reconstruct the relationships between cousins.

To visualize the relationships between wells as determined by the average prefix method, we converted the average prefix to a measure of distance by the following equation:
distance = 1/(average prefix – shortest prefix found in pairing)
Then, we arbitrarily set the shortest distance to 0.5 inches for visualization, and set the other distances proportionally.

All the analyses were done in python. The scripts are available at gitlab.com/__mason__/CHRYON.

### Supplementary Tables (see attached .xls files)

**Table S1.** 293T cells were transfected with plasmids expressing Cas9 alone or Cas9 and TdT, along with one of 16 guide RNAs targeting genomic sites, and allowed to grow for three days, then editing outcomes were determined by NGS. For each length of insertion at each site, the proportions of each nt present in the top (non-target) strand were recorded. The proportions of total insertions of each length were also calculated, in order to determine the average Shannon entropy per bp inserted and the average insertion length over all 16 sites. Full sequences of the target sites and PCR primers used to analyze them are also listed.

**Table S2.** Data underlying the lineage reconstruction in Figure S4A-D. Basic statistics describing the data set are listed, and a full list of insertion sequences for each well is included.

**Table S3.** Data underlying the lineage reconstruction in Figure S4E-F. Basic statistics describing the data set are listed, and a full list of insertion sequences for each well is included.

**Table S4.** Data underlying the lineage reconstruction in Figures 5, S5, and S6. Basic statistics describing the data set are listed, and a full list of insertion sequences for each well is included.

**Table S5.** All data underlying Figure S6.

**Table S6.** Tabulation of all 1- and 2-nt sequences in the data underlying Figures 5, S5, S6, and S7, and Shannon entropy calculations based on these results.

**Table S7.** A guide to plasmids, primers, and NGS data sets produced in this work.

### Supplementary Discussion

*Development of CHYRON components and protocols*

We performed further experiments to test whether our key result shown in Figure 2, that TdT promotes insertions, was correctly interpreted. We showed that Cas9 was expressed at similar levels in the presence or absence of TdT in the experiment shown in Figures 2A, B, and D (**Figure S1A**), and that TdT can promote insertions at all genomic sites tested (**Figure S1B**). To ensure that the reporting of these insertions (which are GC-rich – Figure 2C) is not skewed by our library prep, we tested the relative amplification of synthetic templates with varying proportions of GC nucleotides by two polymerases (**Figure S1C**). Since Q5 polymerase produced a less skewed result, we used it for all library prep PCR steps, except the initial PCR of the CHYRON loci from genomic DNA, for which we found that Phusion was substantially more efficient (data not shown). Finally, we showed that, as expected, dTdT, which lacks residues critical for its polymerase activity[41-42], did not promote insertions at Cas9 cut sites (**Figure S2A**).

Next, we attempted without success to improve the ratio of insertions to deletions at the genomic cut site shown in Figure 2. First, we increased the ratio of TdT to Cas9 transfected (**Figure S2B**). Then we fused TdT to Cas9 through a variety of flexible linkers (**Figure S2C**). We observed TdT activity, but it was reduced relative to constructs in which TdT was expressed without any linkage. Thus, TdT is inhibited, whether sterically or through misfolding, by fusion to Cas9. It may have some activity in the context of a fusion protein, or the activity we see may be due to cleavage of the flexible linker and liberation of TdT (**Figure S2D**).

*Further characterization of $CHYRON_{20}$*

We verified Cas9 expression during the timecourse shown in Figure 3B-D (**Figure S3A**) and included the full plots for Figure 3E-F (**Figure S3B-C**).

*Requirements for successful lineage reconstruction*

In early efforts (data not shown), we attempted to reconstruct relationships among populations of cells using library prep and sequencing protocols that captured an insufficient percentage of the cells in each population. These attempts were unsuccessful because sequencing a high proportion of each population is essential to successful lineage reconstruction. The CHYRON locus from a cell can potentially report on the relationships between the populations in which the cell ends up only if the cell acquires an insertion, then the cell divides one or more times, and then the descendants are split so at least one descendant is distributed to each daughter population. However, the fraction of these potential "reporter" CHYRON loci that will actually give useful information declines with the square of the fraction of each population sampled. Specifically, if r = number of insertions in a data set that arose, then the cell divided, then the daughter cells were split into separate wells and p = proportion of cells in each population that are sampled, then

$$informative\ loci = rp^2$$

For example, consider the following scenario: if only 10% of each well is sampled, the percentage of potential reporter loci that can report accurately is only 0.1 x 0.1 = 0.01, or 1%. If only a small fraction of each population is sampled, the chance of sampling related loci from both of two or more related populations will be very low.

As we attempt to reconstruct population lineage correctly, we must also consider potentially misleading sequences. “Convergent insertions,” or insertions with the same sequence that were generated independently, could cause the unrelated wells in which they arose to be incorrectly considered related. The misleading effect of convergent insertions can be minimized by limiting reconstruction to insertions that are long enough that convergence is unlikely. Assign the following additional variables:

s = average Shannon entropy for insertions of a given length
n = the average number of insertions of that length in a well or population

For an insertion of a given length in well A, the formula for the expected number of convergent insertions in other wells (*i.e.* the expected number of convergent insertions between one of those populations and an unrelated population) is

$$convergent\ insertions = pn\left[1 - \left(\frac{2^s - 1}{2^s}\right)^{pn}\right]$$

We may define ‘effectiveness” as the expected number of matched insertions between two identical populations (*i.e.*, true matches indicating a lineage relationship) divided by the expected number of convergent insertions between one of those populations and an unrelated population.

$$effectiveness = \frac{rp^2}{pn\left[1 - \left(\frac{2^s - 1}{2^s}\right)^{pn}\right]}$$

This can be rewritten as:

$$effectiveness = \frac{rp}{n\left[1 - \left(\frac{2^s - 1}{2^s}\right)^{pn}\right]}$$

Thus, more information-encoding capacity is required as population relatedness and sampling declines, or the population size increases.

*Early attempts at lineage reconstruction*

When we had improved our library prep protocol, we attempted to reconstruct the relationships between populations of cells using $CHYRON_{20}$. In our first attempt (**Figure S4A**), we seeded 10,000 293T-$CHYRON_{20}$ cells, allowed them to double, transfected with Cas9 and TdT, then split each well into two daughter wells each day. After two rounds of splitting, we allowed the cells to grow out for two days, then collected the cells and sequenced the CHYRON loci. With optimized reconstruction parameters, we were able to reconstruct the sibling and cousin relationships of 7 of the 8 wells (**Figure S4B**). There were two parameters we needed to optimize: abundance cutoff and length cutoff. To determine the proper abundance cutoff, we assigned a percentage abundance value to each unique insertion in a well. Then, for each abundance value, we plotted the counts of insertions that had that value (**Figure S4C**). Abundance cutoffs should be set so that, as far as possible, genuine insertion sequences that arise in cells should be included, while those “insertion sequences” that result from library prep errors or sequencing errors should be excluded. If PCR amplification during library prep was relatively even across all sequences, all insertion sequences that arose in cells should be detected in more reads than all artifactual "insertion sequences," because genuine insertion sequences have been amplified through all PCR cycles in the library prep protocol, while artifactual sequences that arose from substitutions introduced during library prep will necessarily have been amplified for fewer cycles. Therefore, when the numbers of insertions having each percentage abundance value are plotted, there should be at least two peaks. The top of the "higher abundance" peak should represent the lowest-abundance genuine insertions and should therefore be the abundance cutoff. Whether due to insufficient sequencing or a low-quality library prep, the cutoff value for this experiment was not clear (**Figure S4C**). Therefore, we attempted reconstructions with a variety of length cutoffs at each possible abundance cutoff (**Figure S4D**). The reconstructions were scored by awarding a maximum of two points for each well: one point for grouping the well with its sibling and one point for grouping the well, at the next level, with at least three of its group of four cousins and sibling. Thus, for this experiment including eight wells, a perfect reconstruction would be awarded a score of 16. For most sets of parameters, the reconstructions were much better than chance, but they fell short of perfect, and there is no clear trend in the parameters that led to better reconstructions (**Figure S4D**). This failure is likely due to inadequate sampling; of long insertions (>9 bp), <10% were shared between sibling wells (**Table S2**). Thus, our sampling efficiency was likely below 10%, and it is not surprising that our reconstructions were not robust.

To further explore the importance of sampling efficiency, we performed a lineage tracing experiment in which we applied a significant bottleneck at each splitting step, and allowed the cells to grow out significantly between each step. We treated 293T-CHYRON$_{20}$ cells with a short pulse of Cas9 and TdT expression just after seeding ~10,000 cells in a 96-well plate, then allowing the cells to double 3-4 times before seeding 1/10 (~8,000) of the cells in each of two wells, briefly pulsing Cas9 and TdT again, then repeating the process (**Figure S4E**). (The cells likely required 9 days to divide 3-4 times after the first transfection because of substantial cell death due to sparse plating.) Using this protocol, we were able to increase our sampling efficiency as expected: >50% of long insertions were shared between sibling wells (**Table S3**). The relationships between the resulting 4 wells were reconstructed perfectly after sequencing the CHYRON locus from each population (**Figure S4F**).

*Further analysis of successful reconstruction*

We used these results to develop a protocol that more-faithfully mimicked a genuine developmental process, without bottlenecking, but could still be reconstructed. As shown in Figure 5, using this new protocol, and the improved CHYRON$_{16i}$, we were able to perfectly reconstruct the relationships of 16 populations including all of the descendants of 40,000 starting cells grown for 10 days. For the purposes of informing future applications of CHYRON (and indeed all DNA recording systems) in complex lineage tracing contexts, we analyzed the interplay between convergence and sampling in greater detail.

In our lineage tracing experiment, reconstruction was aided by different lineages within each population that begin to accumulate insertions at the CHYRON locus at different points in the experiment. To test whether this aid was necessary for our successful reconstruction, we needed to develop a novel reconstruction algorithm that could determine the relatedness of sequences that were not perfectly identical. The "average prefix method" takes advantage of the ordered nature of insertions at the CHYRON locus to determine the relatedness of two wells based on the average prefix of the closest match in the other well of each sequence in each well. The average prefix method accurately reconstructed the splitting procedure (**Figure S5A**). We showed that accumulation of sequences in different cells at different times in the lineage splitting experiment was not crucial to our reconstruction by the average prefix length method: when we computationally exclude all sequences that are identical between cousins, we can still reconstruct the relationships between the populations accurately (**Figure S5B**).

Setting an abundance cutoff properly is essential for accurate reconstructions, lest artifactual "sequences" be included, or too many genuine sequences filtered out. For the experiment shown in Figure 5, for each percentage abundance value, we plotted the counts of insertions that had that value. However, we could not use this plot to determine a proper abundance cutoff because it had only one peak (data not shown). There are two possible contributors to this observation: (1) we did not sequence our library deeply enough (we analyzed ~3M reads per well and each well contained ~1.2M cells with non-root CHYRON loci that should escape PmlI degradation) and/or (2) our library prep process was flawed, so that many sequences that arose as errors during library prep were better-represented in our data set than genuine sequences. To set an abundance cutoff we used a smaller data set that was produced using the same library prep protocol, but was fully sequenced and produced two peaks (**Figure S5B**). Thus, we set an abundance cutoff so that all sequences that were observed in at least 0.0139% of all non-deletion reads in a well were included in the analysis of that well.

In our experiment, we found that >40% of long insertion sequences (unlikely to be identical by chance) were identical between sibling wells (**Table S4**), suggesting that our sampling efficiency is high. Therefore, it is not surprising that our reconstruction is robust: compared to the ideal reconstruction that results from using insertions 8-15 bp in length (Figure 5B), reconstructions using shorter insertions that could arise convergently, or restricted to longer, less abundant insertions only slightly exaggerate the distance between some sibling wells (**Figure S5C**).

By artificially degrading our data set, we were able to systematically test the effect of insertion length cutoffs on reconstruction quality. If a large number of artifactual sequences are included in the data set, because the abundance cutoff is set too low, only reconstructions that rely on longer insertions are perfectly accurate (**Figure S5D**). When insertions were computationally removed at random from each well to simulate a sampling efficiency that is 25% of the actual experiment, there is a trend toward better reconstruction with insertion length cutoffs of 8-15 bp or 9-15 bp (**Figure S5E**). Both of these results can be understood as a reflection of the competition between identical insertion sequences that share a common ancestor and identical insertion sequences that arose independently. The former decline in abundance with the square of the sampling efficiency, as discussed above, so when sampling is reduced to 25%, identical insertion sequences that share a common ancestor are reduced to 6.25% of their former abundance in the pool of insertions sequences used to generate the reconstruction. In contrast, identical insertion sequences that arise independently are only reduced in abundance linearly with declining sampling efficiency, so 25% of these sequences are still present. Longer identical insertions are less likely to arise independently, so reconstructions that rely on longer insertions are more successful when sampling is limited (until the length cutoff is pushed so high that not enough insertions are available to allow reconstruction). Reducing the abundance cutoff introduces artifactual sequences to the data set. These artifactual sequences can be coincidentally identical to sequences in other wells, but of course they cannot reflect true relationships between wells, since they did not arise during the growth of the cells. Therefore, when the abundance cutoff is set too low, setting a longer length cutoff can promote accurate reconstruction by screening out convergent sequences that are coincidentally identical to insertion sequences in unrelated wells. Although it is possible to sample a high proportion of cells in a real-world tissue (Chan et al., 2019), sampling at high efficiency may be challenging in some applications. The high diversity of sequences that can be recorded at the $CHYRON_{16i}$ locus may therefore be crucial for accurate lineage reconstruction.

Finally, we took advantage of the simplicity of the CHYRON output to examine a subset of the data in more detail. We created a list consisting of each insertion sequence in each well. Then, we culled this list so that it includes only 14-bp insertions sharing at least an 8-bp prefix with at least one other insertion, either in the same well or another well (**Table S5**). We alphabetized this list, and then, by hand, highlighted each set of sequences that share an 8-bp prefix (**Figure S7**). These sets are mostly identical sequences shared in sibling wells. Identical sequences or prefixes could also be shared between groups of sibling and cousin wells. Sequences shared between unrelated wells were rare. Just the 14-bp insertions we analyzed (by hand, in a matter of minutes) contained sequences shared between all siblings and all but one set of cousins in the experiment. This simple method of alphabetizing insertions may, for future versions of CHYRON, be sufficient for single-cell-resolution lineage reconstruction.

*Determination of entropy per bp by examining the abundance of each di-nucleotide*

For the data underlying the lineage reconstruction in Figures 5, S5, S6, and S7, we determined the frequencies at which every possible single- and di-nucleotide sequence were observed, and used these frequences to calculate that the average entropy of $CHYRON_{16i}$ in this experiment was 1.9 bits per bp, based on the single nucleotide data, or 1.73 bits per bp, based on the di-nucleotide data (**Table S6**).

### Supplementary Figures

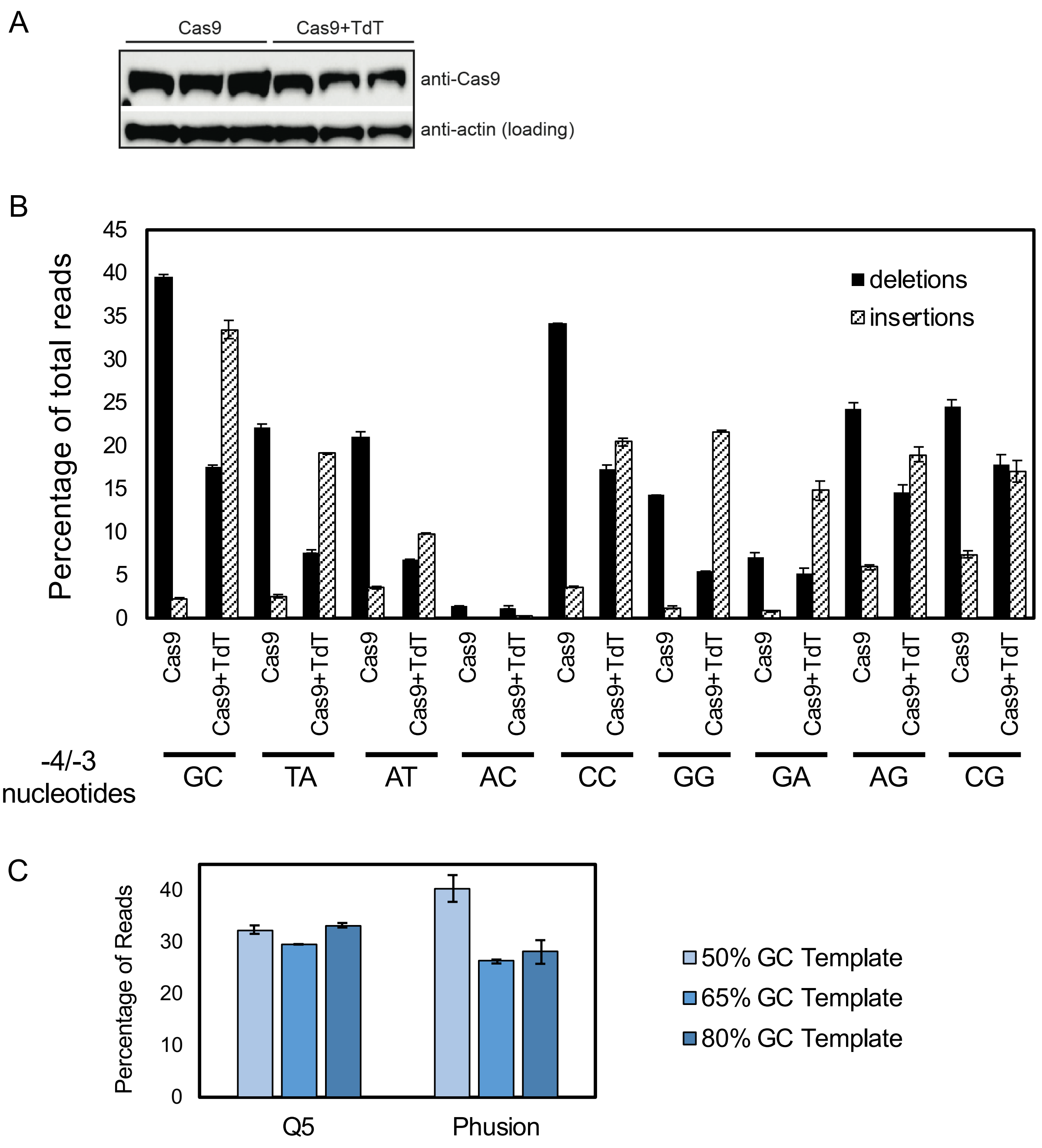


**Insertion sequences for the 50%, 65%, and 80% GC-rich templates for the PCR bias assay of Q5 and Phusion polymerases.**

| % GC | 5'-Insertion Sequence-3' |
|---|---|
| 50 | CATGGCATGGATCCTGTGACTCGATATAGGAACTCCGATC |
| 65 | CCGTGTCAGCTCACTGCGATCCAGGGTCCACTCCCAGTGC |
| 80 | CCGTGGCAGCTCGCTGCGCCAGGGTCCACGCCCAGGGC |

**Figure S1.**

**Figure S1. Expression of Cas9 upon transfection, effect of TdT on editing outcomes at a variety of genomic sites, and test of fidelity of NGS library prep for GC-rich templates. Related to Figure 2 and Methods. (A)** Western blot of samples from Figure 2A-B. **(B)** Percentages of deletions and insertions at different -4/-3 nucleotide cut sites. HEK293T cells were transfected with the appropriate sgRNA and either a Cas9 or Cas9-T2A-TdT construct. Bars represent the average and error bars represent the range of two biological replicates. **(C)** Percentages of the total number of amplicons from a 1:1:1 molar ratio of the 50%, 65%, and 80% GC-rich templates. Error bars represent the range of two technical replicates.

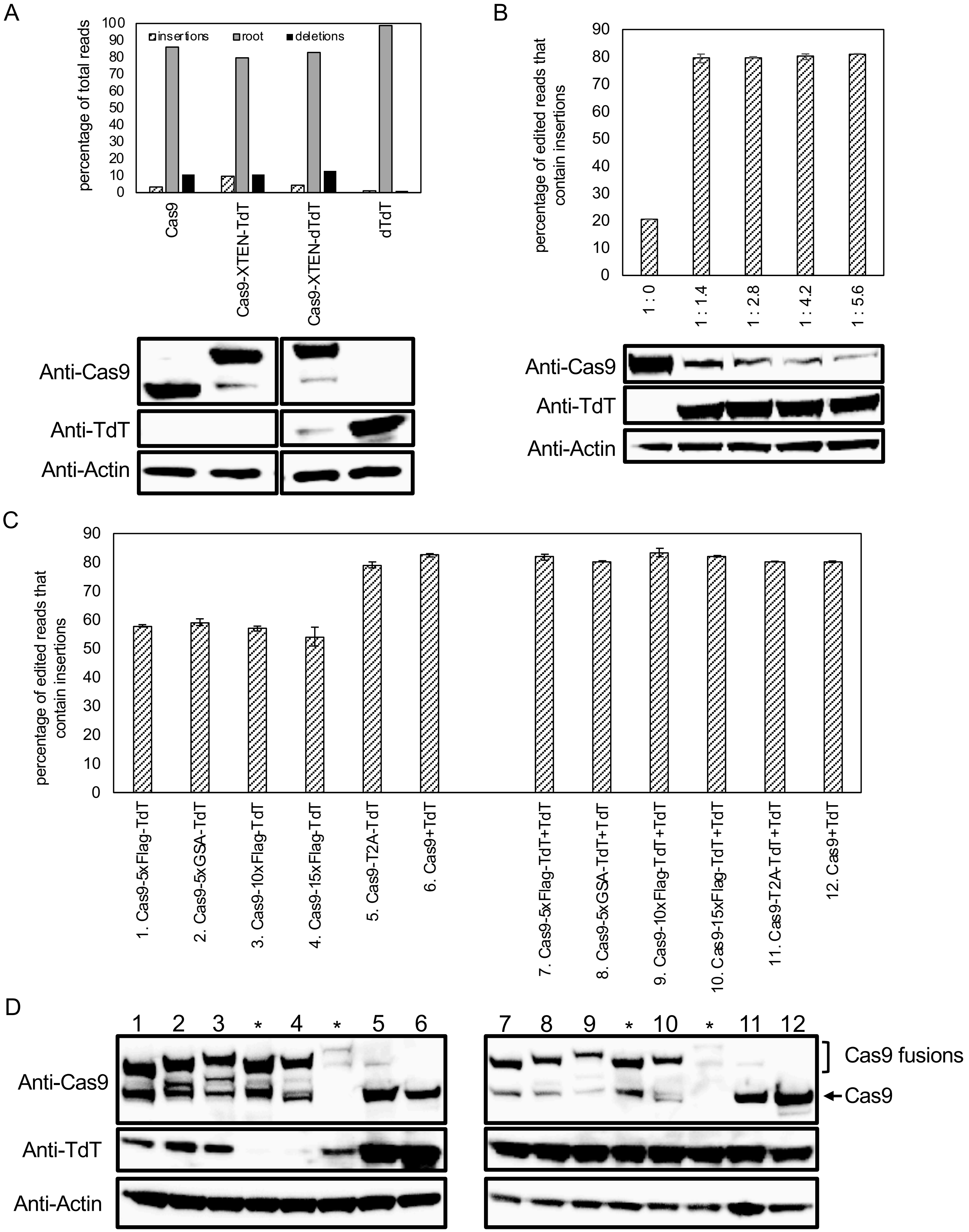
A
percentage of total reads
insertions
root
deletions
Cas9
Cas9-XTEN-TdT
Cas9-XTEN-dTdT
dTdT
Anti-Cas9
Anti-TdT
Anti-Actin
B
percentage of edited reads that contain insertions
1 : 0
1 : 1.4
1 : 2.8
1 : 4.2
1 : 5.6
Anti-Cas9
Anti-TdT
Anti-Actin
C
percentage of edited reads that contain insertions
1. Cas9-5xFlag-TdT
2. Cas9-5xGSA-TdT
3. Cas9-10xFlag-TdT
4. Cas9-15xFlag-TdT
5. Cas9-T2A-TdT
6. Cas9+TdT
7. Cas9-5xFlag-TdT+TdT
8. Cas9-5xGSA-TdT+TdT
9. Cas9-10xFlag-TdT+TdT
10. Cas9-15xFlag-TdT+TdT
11. Cas9-T2A-TdT+TdT
12. Cas9+TdT
D
1 2 3 * 4 * 5 6
7 8 9 * 10 * 11 12
Anti-Cas9
Anti-TdT
Anti-Actin
Cas9 fusions
Cas9

**Figure S2.**

**Figure S2. Activity of varying amounts and fusions of TdT. Related to Figure 2. (A)** TdT promoted increased insertions at Cas9 cut sites via its polymerase activity. A catalytically dead version of TdT, “dTdT,” was created by introducing the mutations D343E and D345E. 293T cells were transfected with plasmids expressing Cas9, Cas9-XTEN-TdT, Cas9-XTEN-dTdT, or dTdT alone, and an sgRNA against a genomic site (HEK293site3). Three days later, cells were collected, DNA was extracted, and the targeted genomic site was amplified by PCR and sequenced by NGS. Sequences were annotated as unchanged (root); pure insertions (insertions); or any sequence that leads to a loss of information (deletions). Western blots were performed on equal amounts of protein from each sample. **(B)** An increased ratio of TdT to Cas9 near our current range did not increase insertion rate. Plasmids expressing Cas9 and TdT were mixed in the ratios noted and transfected into 293T cells along with an sgRNA against a genomic site (HEK293site3). Three days later, cells were collected, DNA was extracted, and the targeted genomic site was amplified by PCR and sequenced by NGS. Sequences were annotated as unchanged, or changed. Then, the proportion of changed sequences that were pure insertions was calculated. Bars represent the average and error bars represent the range of two technical replicates. Western blots were performed on equal amounts of protein from each sample. **(C)** Expressing Cas9-TdT fusions promoted insertions, but less efficiently than free TdT. Plasmids expressing various Cas9-TdT fusions were transfected into 293T cells along with an sgRNA against a genomic site (HEK293site3), with or without additional free TdT. Three days later, cells were collected, DNA was extracted, and the targeted genomic site was amplified by PCR and sequenced by NGS. Sequences were annotated as unchanged, or changed. Then, the proportion of changed sequences that were pure insertions was calculated. Bars represent the average and error bars represent the range of two technical replicates. **(D)** Western blots were performed on equal amounts of protein from each sample shown in (C). * represents fusion proteins not included in the analysis.

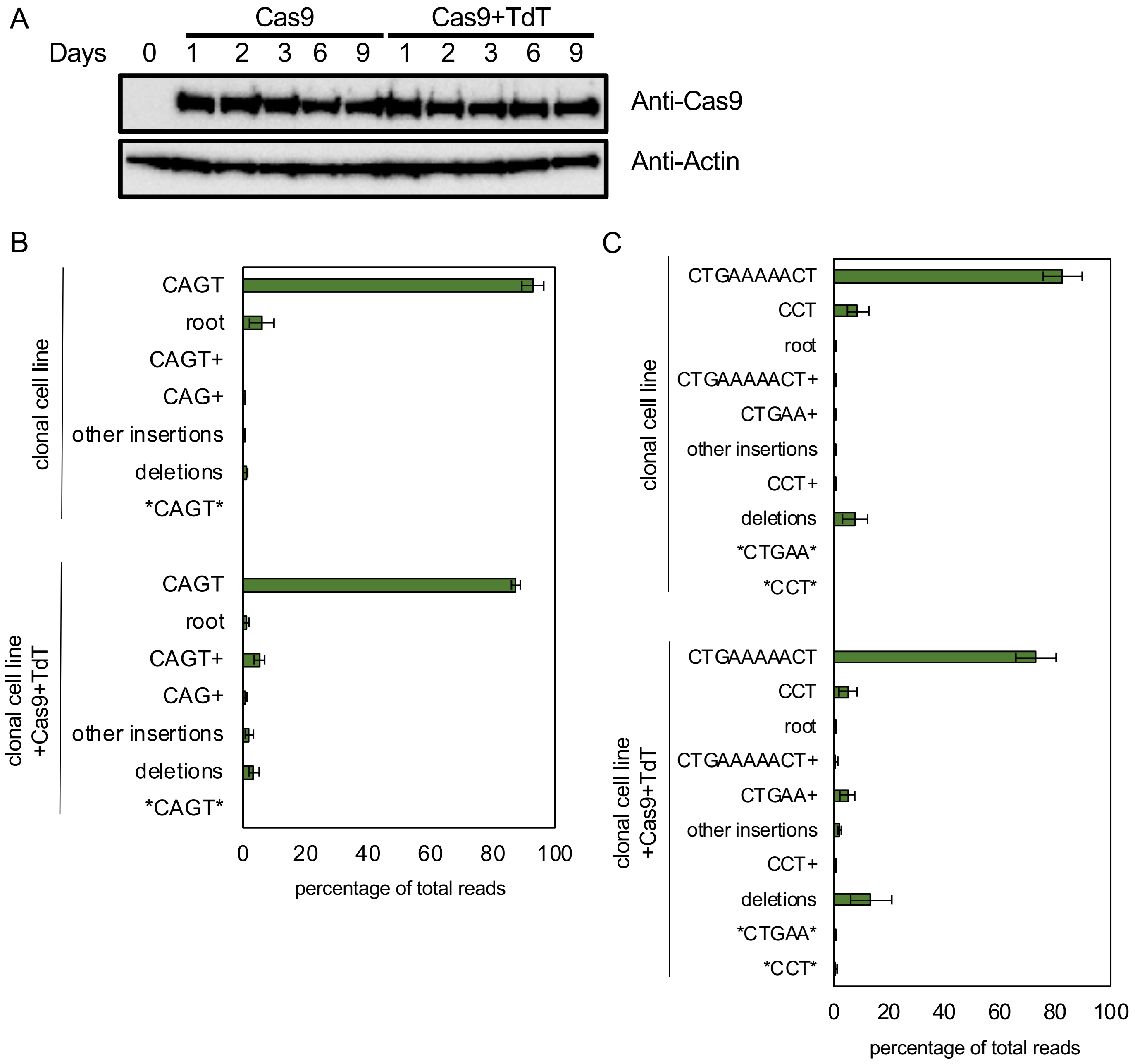
A
Cas9
Cas9+TdT
Days 0 1 2 3 6 9 1 2 3 6 9
Anti-Cas9
Anti-Actin
B
clonal cell line
CAGT
root
CAGT+
CAG+
other insertions
deletions
*CAGT*
clonal cell line +Cas9+TdT
CAGT
root
CAGT+
CAG+
other insertions
deletions
*CAGT*
0 20 40 60 80 100
percentage of total reads
C
clonal cell line
CTGAAAAACT
CCT
root
CTGAAAAACT+
CTGAA+
other insertions
CCT+
deletions
*CTGAA*
*CCT*
clonal cell line +Cas9+TdT
CTGAAAAACT
CCT
root
CTGAAAAACT+
CTGAA+
other insertions
CCT+
deletions
*CTGAA*
*CCT*
0 20 40 60 80 100
percentage of total reads

**Figure S3.**

**Figure S3. Further characterization of $CHYRON_{20}$. Related to Figure 3. (A)** Western blots of samples shown in Figure 3B-C. **(B)** Cas9 and TdT mediated multiple rounds of editing on an integrated hgRNA. The 293T-$CHYRON_{20}$ cell line was transfected with Cas9 and TdT to induce insertions, then purified to near-clonality. This near-clonal cell line bearing an insertion with the sequence CAGT was then transfected again with a plasmid expressing Cas9 and TdT. These cells, and an untransfected control, were grown for 6-15 days, then collected. The CHYRON locus was sequenced and editing outcomes were determined to be the root $CHYRON_{20}$ sequence (root), deletions, a CAGT insertion (CAGT), an insertion containing the prefix CAGT or CAG (CAGT+ or CAG+, respectively), an insertion containing the sequence CAGT other than as a prefix (*CAGT*), or other insertions. Error bars=±stdev of two technical replicates each of the 6 and 15 day timepoints. This is the full plot from Figure 3E. **(C)** Another near-clonal cell line was isolated, this one bearing the insertions CTGAAAAACT and CCT. This cell line was treated as in (D) and the CHYRON locus was sequenced and editing outcomes were determined to be the root $CHYRON_{20}$ sequence (root), deletions, both isolated insertions (CTGAAAAACT and CCT), an insertion containing these insertions or a shortened version as a prefix (CTGAAAAACT+, CCT+, CTGAA+), an insertion containing the sequences CTGAA or CCT other than as a prefix (*CTGAA* and *CCT*), or other insertions. Error bars=±stdev of two technical replicates each of the 6 and 15 day timepoints. This is the full plot from Figure 3F.

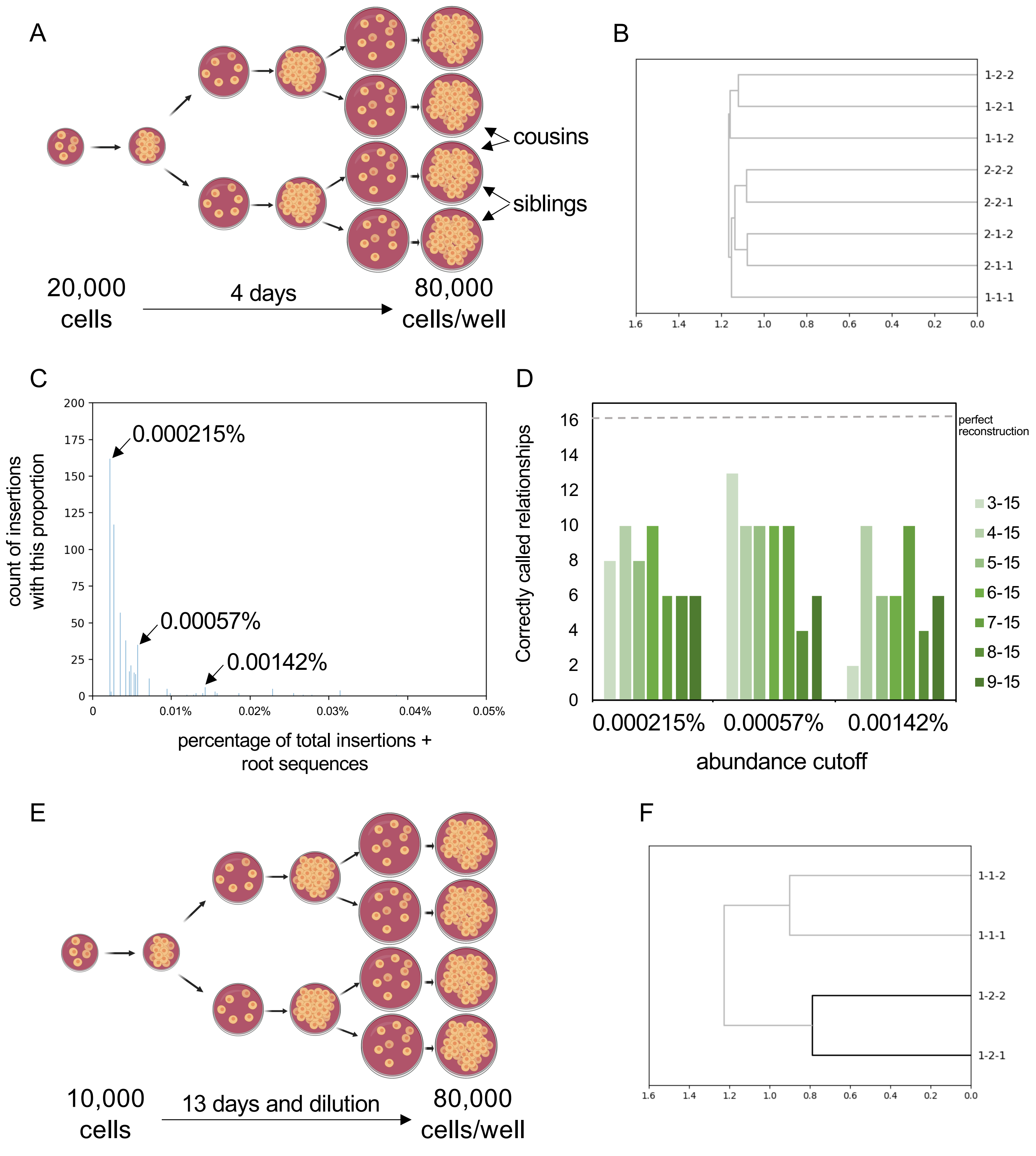
A
cousins
siblings
20,000
cells
4 days
80,000
cells/well
B
1-2-2
1-2-1
1-1-2
2-2-2
2-2-1
2-1-2
2-1-1
1-1-1
1.6 1.4 1.2 1.0 0.8 0.6 0.4 0.2 0.0
C
count of insertions with this proportion
0.000215%
0.00057%
0.00142%
200 175 150 125 100 75 50 25 0
0 0.01% 0.02% 0.03% 0.04% 0.05%
percentage of total insertions + root sequences
D
Correctly called relationships
perfect reconstruction
16 14 12 10 8 6 4 2 0
3-15
4-15
5-15
6-15
7-15
8-15
9-15
0.000215% 0.00057% 0.00142%
abundance cutoff
E
10,000
cells
13 days and dilution
80,000
cells/well
F
1-1-2
1-1-1
1-2-2
1-2-1
1.6 1.4 1.2 1.0 0.8 0.6 0.4 0.2 0.0

**Figure S4.**

**Figure S4. Robust lineage reconstruction with CHYRON$_{20}$ is only possible for splitting protocols with good sampling efficiency. Related to Figure 5. (A)** Plan for splitting in which 293T-CHYRON$_{20}$ cells are split approximately once per cell division. **(B)** 7 of 8 wells were properly placed in a lineage tree for a narrow range of reconstruction parameters. Lineage reconstruction using all 3-15 bp insertions that represent greater than 0.00057% of all insertion or root sequences to calculate Jaccard similarity followed by UPGMA hierarchical clustering. **(C)** The best count cutoff for the experiment shown in (A) and (B) was unclear. The total number of reads in each well bearing either the root sequence or an insertion was calculated. Then, the number of reads for each unique insertion sequence in the well was determined and divided by the total to give the percentage abundance of that insertion. For each percentage abundance, the number of insertions with that abundance was plotted. The proper abundance cutoff is that at the top of the higher-abundance peak. The possible abundance cutoffs for these experiments are highlighted. **(D)** Reconstruction was marginal over a variety of parameters. The reconstruction was repeated with different abundance cutoffs and length cutoffs for the insertions used for reconstruction. The quality of the reconstruction was scored. For each well, the reconstruction got one point for grouping the well with the proper sibling well and one point for grouping the well with at least 2 of the 3 siblings and cousins. Thus, a maximum of 16 points are possible for a reconstruction of the relationships of 8 wells. **(E)** Plan for splitting with substantial dilution and grow-out to improve sampling efficiency. **(F)** This splitting protocol allowed perfect lineage reconstruction. Lineage reconstruction using 9-15 bp insertions to calculate Jaccard similarity followed by UPGMA hierarchical clustering.

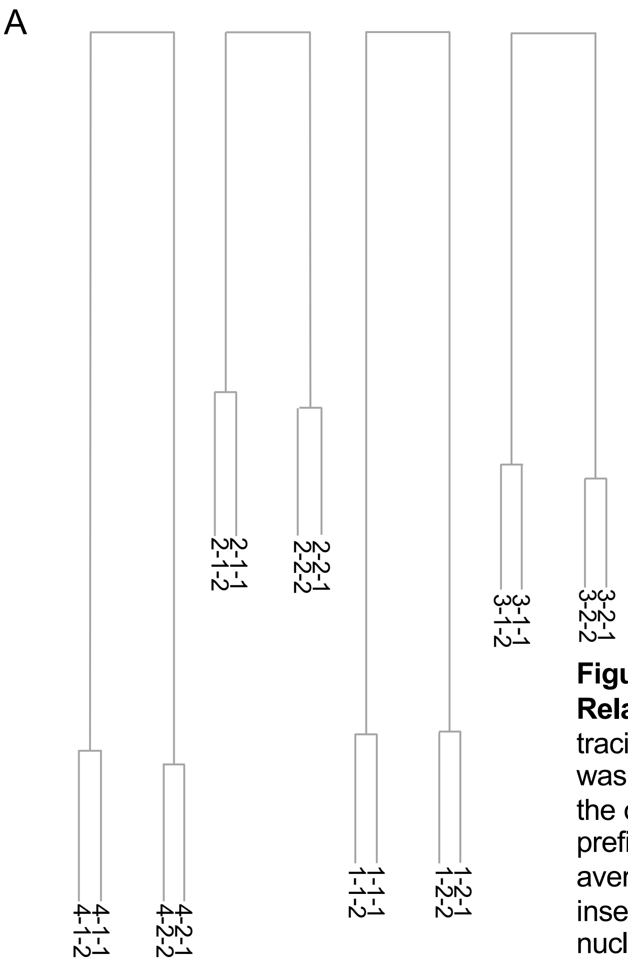


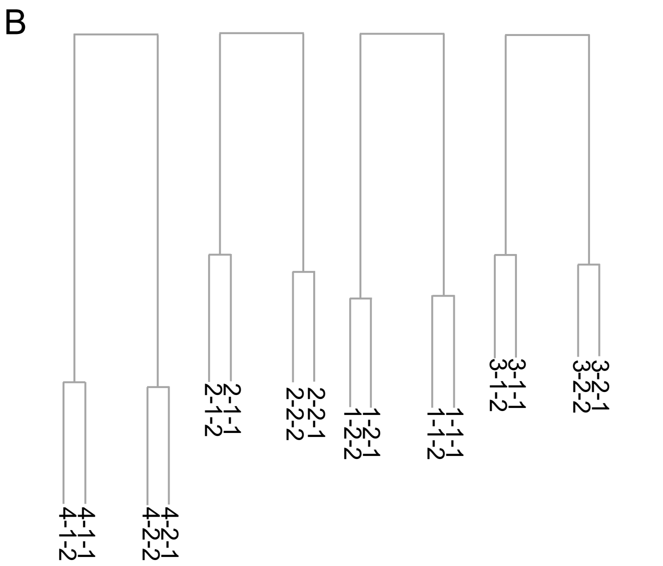


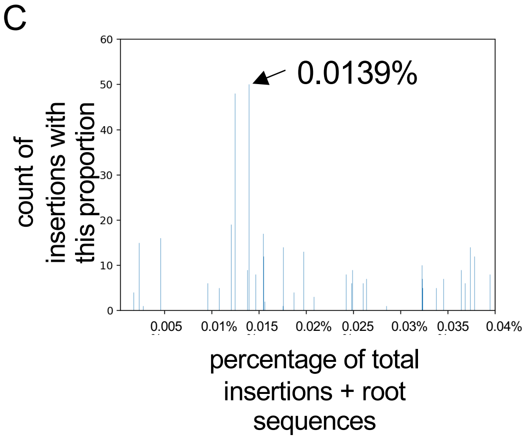


**Figure S5. Lineage reconstruction using $CHYRON_{16i}$ is robust. Related to Figure 5.** This Figure contains analyses of the lineage tracing data set shown in Figure 5. (**A**) The relatedness of all wells was perfectly reconstructed using a new method that makes use of the ordered nature of insertions at the CHYRON locus. An "average prefix" was calculated for all possible pairs of wells. To calculate the average prefix between wells A and B, for example, for each insertion in well A, the insertion in well B with the most initial nucleotides in common (the "longest prefix") is found. For that pairing, the number of initial nucleotides shared between the two sequences (the "prefix length") is recorded. This process is repeated for every other insertion in well A, then for every insertion in well B. The prefix lengths of all of the paired insertions are then averaged to calculate the "average prefix" between the two wells. Once the average prefix for each pair of wells has been calculated, the two wells with the longest average prefix are paired, then the remaining wells with the longest average prefix are paired, and so on until all wells are paired. Thus, all pairs of siblings are identified. Then, for each pair of siblings, a "pooled well" was created, consisting of insertions common to each well. The average prefix process was then repeated on these pooled wells. The distance between each pair of wells was visualized by determining the average prefix of insertions in the two wells, subtracting the average prefix of insertions in the two most-distantly related wells, then inverting the result and setting all the distances proportionally. **(B)** Lineage reconstruction could be performed using insertions that expanded throughout the experiment. All insertions that were identical between a well and either of its cousin wells were computationally removed from the data set. Then, well relationships were reconstructed by the average prefix method, as in Figure 5B, using insertions 8-15 bp in length. The distances shown here are proportional to those in (A). **(C)** The abundance cutoff for this experiment was determined using a smaller, deeply-sequenced dataset that was obtained using the same library prep protocol. The number of insertions with each abundance were plotted as in Figure S4C, and 0.0139% was chosen as the cutoff.

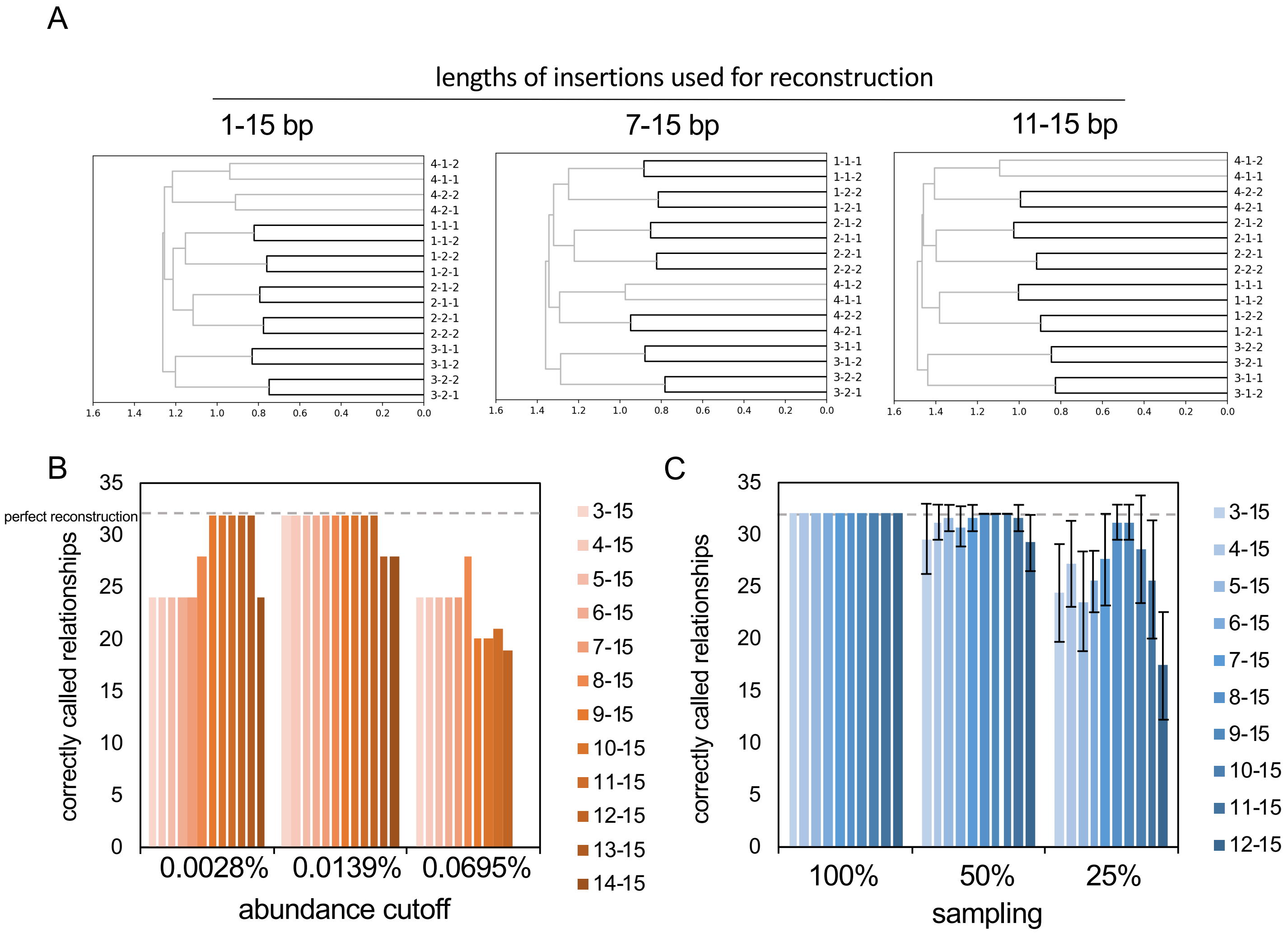


**Figure S6. High-information recording and optimal analysis is required when sampling is limited. Related to Figure 5.** This Figure contains analyses of the lineage tracing data set shown in Figure 5. **(A)** Lineage reconstruction changed little as length cutoffs were changed. Reconstruction was performed by determining Jaccard similarities between pairs of wells and then performing UPGMA hierarchical clustering, as in Figure 5C. Insertions of the indicated lengths were used for reconstruction. **(B)** Abundance cutoffs affected reconstruction. Lineage reconstructions were performed by determining Jaccard distances between pairs of wells and then performing UPGMA hierarchical clustering, using insertions within the abundance and length cutoffs indicated. Reconstructions were scored as in Figure S4D. For each well, the reconstruction got one point for grouping the well with the proper sibling well and one point for grouping the well with at least 2 of the 3 siblings and cousins. Because the relationships between 16 wells were reconstructed, a maximum of 32 points is possible. Bars are missing for the most-stringent combinations of length and abundance cutoffs because there were not enough insertions above the cutoffs to calculate the Jaccard similarity. **(C)** Length cutoffs became more important as sampling efficiency declined. 50% or 75% of insertions were computationally removed at random from the dataset, then lineage reconstruction and reconstruction scoring were performed as in (A). Computational random insertion removal and subsequent reconstruction was performed 10 times for each insertion length cutoff and sampling efficiency. The average reconstruction score is shown (error bars=±stdev).

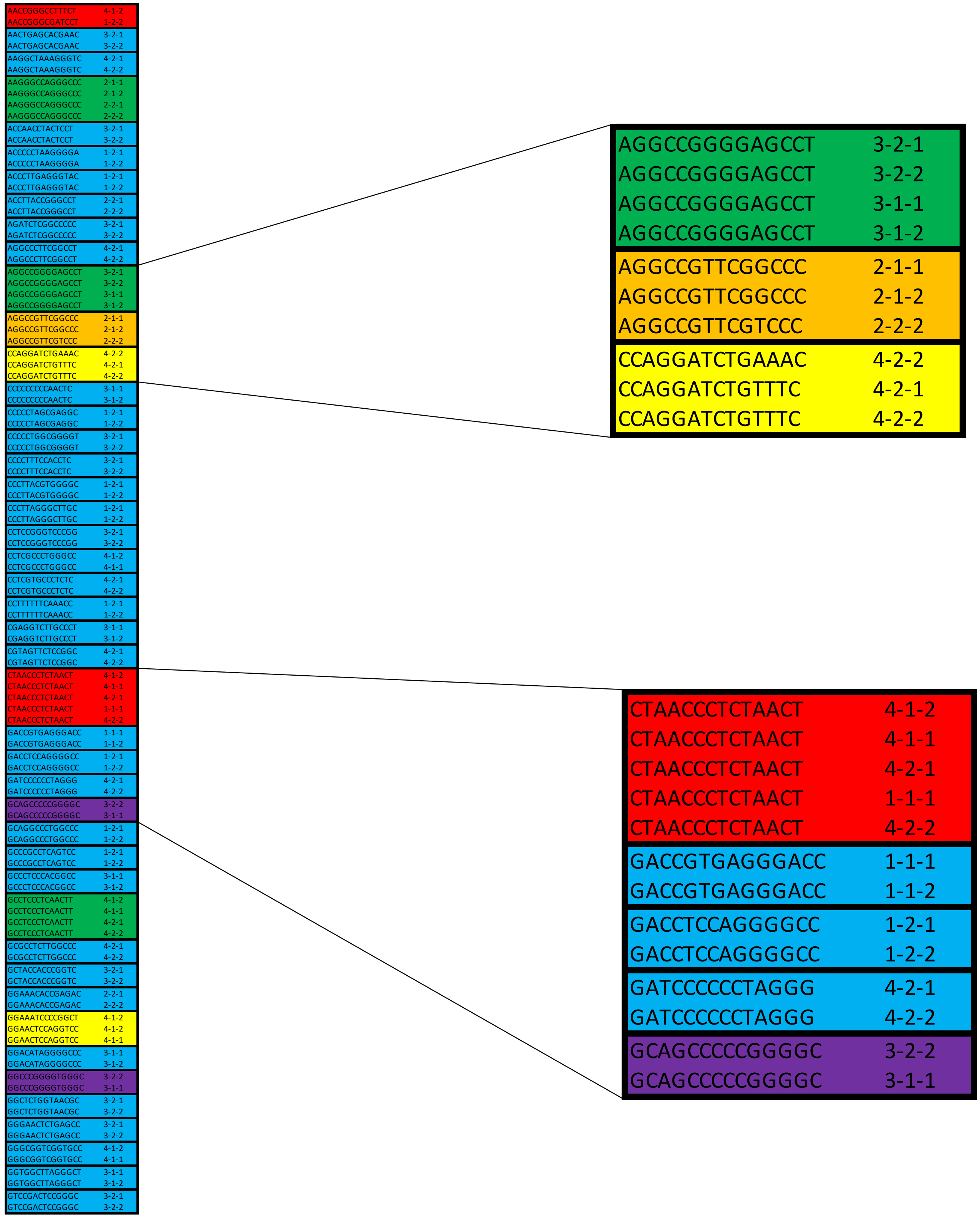
AACCGGGCCTTTCT 4-1-2
AACCGGGCGATCCT 1-2-2
AACTGAGCACGAAC 3-2-1
AACTGAGCACGAAC 3-2-2
AAGGCTAAAGGGTC 4-2-1
AAGGCTAAAGGGTC 4-2-2
AAGGGCCAGGGCCC 2-1-1
AAGGGCCAGGGCCC 2-1-2
AAGGGCCAGGGCCC 2-2-1
AAGGGCCAGGGCCC 2-2-2
ACCAACCTACTCCT 3-2-1
ACCAACCTACTCCT 3-2-2
ACCCCCTAAGGGGA 1-2-1
ACCCCCTAAGGGGA 1-2-2
ACCCTTGAGGGTAC 1-2-1
ACCCTTGAGGGTAC 1-2-2
ACCTTACCGGGCCT 2-2-1
ACCTTACCGGGCCT 2-2-2
AGATCTCGGCCCCC 3-2-1
AGATCTCGGCCCCC 3-2-2
AGGCCCTTCGGCCT 4-2-1
AGGCCCTTCGGCCT 4-2-2
AGGCCGGGGAGCCT 3-2-1
AGGCCGGGGAGCCT 3-2-2
AGGCCGGGGAGCCT 3-1-1
AGGCCGGGGAGCCT 3-1-2
AGGCCGTTCGGCCC 2-1-1
AGGCCGTTCGGCCC 2-1-2
AGGCCGTTCGTCCC 2-2-2
CCAGGATCTGAAAC 4-2-2
CCAGGATCTGTTTC 4-2-1
CCAGGATCTGTTTC 4-2-2
CCCCCCCCCAACTC 3-1-1
CCCCCCCCCAACTC 3-1-2
CCCCCTAGCGAGGC 1-2-1
CCCCCTAGCGAGGC 1-2-2
CCCCCTGGCGGGGT 3-2-1
CCCCCTGGCGGGGT 3-2-2
CCCCTTTCCACCTC 3-2-1
CCCCTTTCCACCTC 3-2-2
CCCTTACGTGGGGC 1-2-1
CCCTTACGTGGGGC 1-2-2
CCCTTAGGGCTTGC 1-2-1
CCCTTAGGGCTTGC 1-2-2
CCTCCGGGTCCCGG 3-2-1
CCTCCGGGTCCCGG 3-2-2
CCTCGCCCTGGGCC 4-1-2
CCTCGCCCTGGGCC 4-1-1
CCTCGTGCCCTCTC 4-2-1
CCTCGTGCCCTCTC 4-2-2
CCTTTTTTCAAACC 1-2-1
CCTTTTTTCAAACC 1-2-2
CGAGGTCTTGCCCT 3-1-1
CGAGGTCTTGCCCT 3-1-2
CGTAGTTCTCCGGC 4-2-1
CGTAGTTCTCCGGC 4-2-2
CTAACCCTCTAACT 4-1-2
CTAACCCTCTAACT 4-1-1
CTAACCCTCTAACT 4-2-1
CTAACCCTCTAACT 1-1-1
CTAACCCTCTAACT 4-2-2
GACCGTGAGGGACC 1-1-1
GACCGTGAGGGACC 1-1-2
GACCTCCAGGGGCC 1-2-1
GACCTCCAGGGGCC 1-2-2
GATCCCCCCTAGGG 4-2-1
GATCCCCCCTAGGG 4-2-2
GCAGCCCCCGGGGC 3-2-2
GCAGCCCCCGGGGC 3-1-1
GCAGGCCCTGGCCC 1-2-1
GCAGGCCCTGGCCC 1-2-2
GCCCGCCTCAGTCC 1-2-1
GCCCGCCTCAGTCC 1-2-2
GCCCTCCCACGGCC 3-1-1
GCCCTCCCACGGCC 3-1-2
GCCTCCCTCAACTT 4-1-2
GCCTCCCTCAACTT 4-1-1
GCCTCCCTCAACTT 4-2-1
GCCTCCCTCAACTT 4-2-2
GCGCCTCTTGGCCC 4-2-1
GCGCCTCTTGGCCC 4-2-2
GCTACCACCCGGTC 3-2-1
GCTACCACCCGGTC 3-2-2
GGAAACACCGAGAC 2-2-1
GGAAACACCGAGAC 2-2-2
GGAAATCCCCGGCT 4-1-2
GGAACTCCAGGTCC 4-1-2
GGAACTCCAGGTCC 4-1-1
GGACATAGGGGCCC 3-1-1
GGACATAGGGGCCC 3-1-2
GGCCCGGGGTGGGC 3-2-2
GGCCCGGGGTGGGC 3-1-1
GGCTCTGGTAACGC 3-2-1
GGCTCTGGTAACGC 3-2-2
GGGAACTCTGAGCC 3-2-1
GGGAACTCTGAGCC 3-2-2
GGGCGGTCGGTGCC 4-1-2
GGGCGGTCGGTGCC 4-1-1
GGTGGCTTAGGGCT 3-1-1
GGTGGCTTAGGGCT 3-1-2
GTCCGACTCCGGGC 3-2-1
GTCCGACTCCGGGC 3-2-2
AGGCCGGGGAGCCT 3-2-1
AGGCCGGGGAGCCT 3-2-2
AGGCCGGGGAGCCT 3-1-1
AGGCCGGGGAGCCT 3-1-2
AGGCCGTTCGGCCC 2-1-1
AGGCCGTTCGGCCC 2-1-2
AGGCCGTTCGTCCC 2-2-2
CCAGGATCTGAAAC 4-2-2
CCAGGATCTGTTTC 4-2-1
CCAGGATCTGTTTC 4-2-2
CTAACCCTCTAACT 4-1-2
CTAACCCTCTAACT 4-1-1
CTAACCCTCTAACT 4-2-1
CTAACCCTCTAACT 1-1-1
CTAACCCTCTAACT 4-2-2
GACCGTGAGGGACC 1-1-1
GACCGTGAGGGACC 1-1-2
GACCTCCAGGGGCC 1-2-1
GACCTCCAGGGGCC 1-2-2
GATCCCCCCTAGGG 4-2-1
GATCCCCCCTAGGG 4-2-2
GCAGCCCCCGGGGC 3-2-2
GCAGCCCCCGGGGC 3-1-1

**Figure S7.**

**Figure S7. Analysis by alphabetizing of insertions at the CHYRON$_{16i}$ locus. Related to Figure 5.** From the data set shown in Figure 5, all 14 bp insertions that share at least the same 8 first letters (an "8 bp prefix") with an insertion in another well or another insertion in the same well were alphabetized in Microsoft Excel. Each group of insertions (marked by their well location) that were identical or shared a prefix greater than or equal to 8 bp was bounded by a black box. The insertions were colored based on their relationship to each other: identical insertions shared only between sibling wells were colored blue, identical insertions shared between all four cousins are colored green, insertions that share a prefix between cousins were colored orange, insertions that share a prefix between siblings were colored yellow, identical insertions shared between less than four cousins were colored purple, and insertions shared between unrelated wells were colored red.
